# Quality of vaccination-induced T cell responses is conveyed by polyclonality and high, but not maximum, antigen receptor avidity

**DOI:** 10.1101/2024.10.30.620795

**Authors:** Katharina Kocher, Felix Drost, Abel Mekonnen Tesfaye, Carolin Moosmann, Christine Schülein, Myriam Grotz, Elvira D’Ippolito, Frederik Graw, Bernd Spriewald, Dirk H. Busch, Christian Bogdan, Matthias Tenbusch, Benjamin Schubert, Kilian Schober

## Abstract

While the quantity of vaccination-induced T cells represents a routine immunogenicity parameter, the quality of such responses is poorly understood. Here, we report on a clinical cohort of 29 human healthy individuals who received three mRNA vaccinations against SARS-CoV-2 before any breakthrough infection. We characterized the magnitude, phenotype and clonal composition of CD8 T cell responses against 16 epitope specificities by ELISpot, flow cytometry as well as single-cell RNA, TCR and surface protein sequencing. To test the functionality of identified clonotypes, 106 T cell receptors (TCR) from five epitope-specific repertoires were re-expressed and tested for peptide sensitivity. While recruited repertoires were overall enriched for high-avidity TCRs, differential clonal expansion was not linked to fine avidity differences. Instead, maintenance of polyclonality ensured robustness in counteracting mutational escape of epitopes. Our findings on the induction and maintenance of high-functionality polyclonal T cell repertoires shed light on T cell quality as a neglected criterion in the assessment of vaccine immunogenicity.

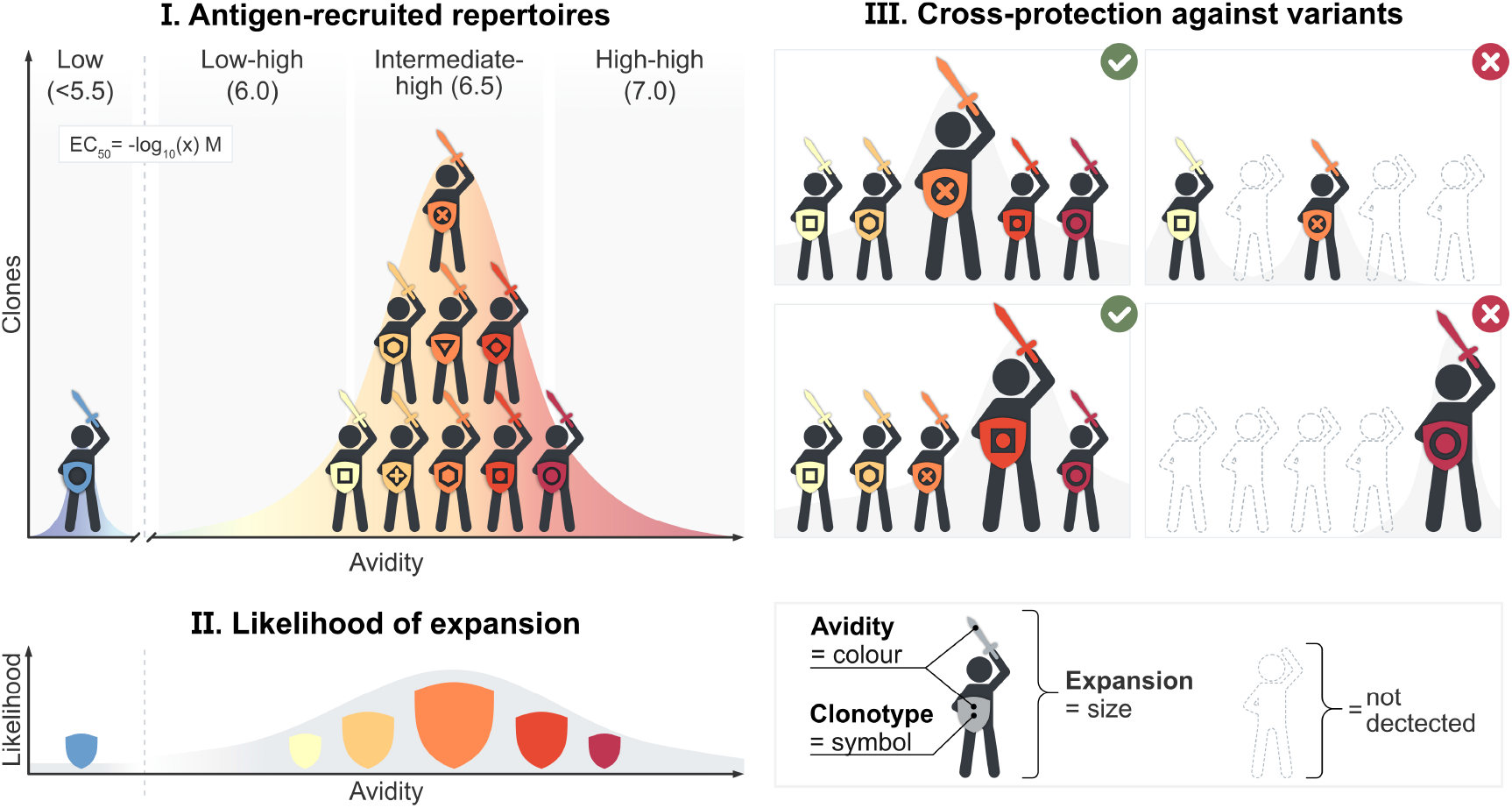

## Introduction

Evaluating immune response quality is essential for the assessment of vaccine immunogenicity and efficacy. The soluble effector molecules of B cells – antibodies – are often characterized in-depth after infection or vaccination. In fact, the neutralization capacity of antibodies is an important correlate of protection for many vaccines^1^. Antibody affinity also serves to indicate the quality of immunogenicity after vaccination^2,3^, and is even being used on a routine basis in the clinics to estimate the timing of *Toxoplasma gondii* exposure in pregnant women^4^. In stark contrast, our understanding of vaccination-induced T cell responses remains superficial. It is generally accepted that T cells are an essential component of long-lasting immunological memory. Yet, for the evaluation of vaccine immunogenicity, T cell responses are often only characterized on a quantitative, but not in-depth on a qualitative level^5^.

T cell quality may refer to the cellular phenotype or metabolic state. While these features are potentially transient, a hallmark of adaptive immunity is the recruitment and expansion of antigen-specific clonotypes with fixed unique antigen receptors^6,7^. The specificity and binding strength (avidity) of a T cell receptor (TCR) is thereby thought to be a major determinant of the expansion, differentiation and protective efficacy of T cell clonotypes^8^. In mouse models of immunization or acute infection, high TCR avidity has generally been shown to correlate with clonal expansion^9–14^. However, during chronic antigen exposure through viral infection or tumor disease, low-avidity clonotypes may also dominate^15,16^.

In humans, dominating antigen-specific T cell populations have similarly been shown to harbor low or high TCR avidities depending on the individual setting^17,18^. Often, such analyses suffer from a scarcity of defined cohorts of healthy individuals who received the same immunizations in synchronized time spans without having encountered the same antigen before. The severe acute respiratory syndrome coronavirus 2 (SARS-CoV-2) vaccination campaign provided a rare research opportunity to follow adaptive immune responses in humans in a systematic manner. Numerous studies have investigated antigen-specific CD8 T cell responses following coronavirus disease 2019 (Covid-19) infection and/or SARS-CoV-2 vaccination^19,20,29–38,21,39,22–28^. These reports provided important insights into the magnitude, phenotype and partly also clonal development of human antigen-specific CD8 T cell responses following vaccination, but did not address whether or how TCR avidity drives clonal repertoire evolution.

Recent findings indicate that adenovirus 5-based human immunodeficiency virus (HIV) vaccines induce CD8 T cells with low TCR avidity and that this could explain a lack in clinical protection^40^. It remains, however, technically challenging to measure the avidity of human antigen-specific T cell responses in a comprehensive manner^41,42^. Which clonotype-intrinsic T cell qualities are induced by vaccination, and how clonal expansion is balanced out with maintenance of diverse repertoires, is therefore still unclear.

## Results

### Identification of SARS-CoV-2 antigen-specific CD8 T cell responses

Within the “CoVa-Adapt” study, we analyzed a cohort of 29 health-care workers who received three doses of the mRNA vaccine Comirnaty (**Table 1; Suppl. Table 1**). Peripheral blood mononuclear cells (PBMCs) and serum were sampled in the acute phase, 10 days after each vaccination (P10, S10, T10) (**Fig. 1A**). Additional samples were collected at early and late memory time points after the 2^nd^ (S68, S210) and 3^rd^ dose (T108, T189). All donors were seronegative for SARS-CoV-2 at the time of the first immunization and did not experience a breakthrough infection until after T10. Antibodies and T cells were first analyzed on a quantitative level. As previously described^43^, all donors had detectable spike-specific IgG levels at S10 that underwent contraction at S210, were boosted at T10, and were maintained later on (T189) especially when an additional breakthrough infection occurred (**Fig. 1B**). Spike-specific IgA and IgM were most prominent at S10 (**Suppl. Fig. 1A**). Frequencies of spike-reactive T cells, determined by *in vitro* re-stimulation with SARS-CoV-2 wild type strain spike antigen, also peaked at S10 in all donors and were similarly boosted by the 3^rd^ vaccination as shown by ELISpot and flow cytometric intracellular cytokine analysis (**Fig. 1C-D, Suppl. Fig. 1B-C**). Spike-reactive CD8 T cells were detected at lower and more heterogenous frequencies, likely due to less efficient re-stimulation with untrimmed 15-mer peptides (**Fig. 1D, Suppl. Fig. 1B-C**). As previously described^44^, spike-reactive T cells persisted months after vaccination. Some spike-reactivity was observed prior to vaccination, likely due to cross-reactivity towards common cold coronaviruses^34,45^. While spike-specific IgG antibodies only mildly correlated with spike-reactive CD8 T cell frequencies (r: 0.36), we observed a stronger correlation with CD4 T cells (r: 0.75), demonstrating the close link between these two pillars of the immune system (**Suppl. Fig. 1D**). Overall, a stable and polyfunctional immune response was established against SARS-CoV-2 spike in all analyzed donors.

**Table 1:**
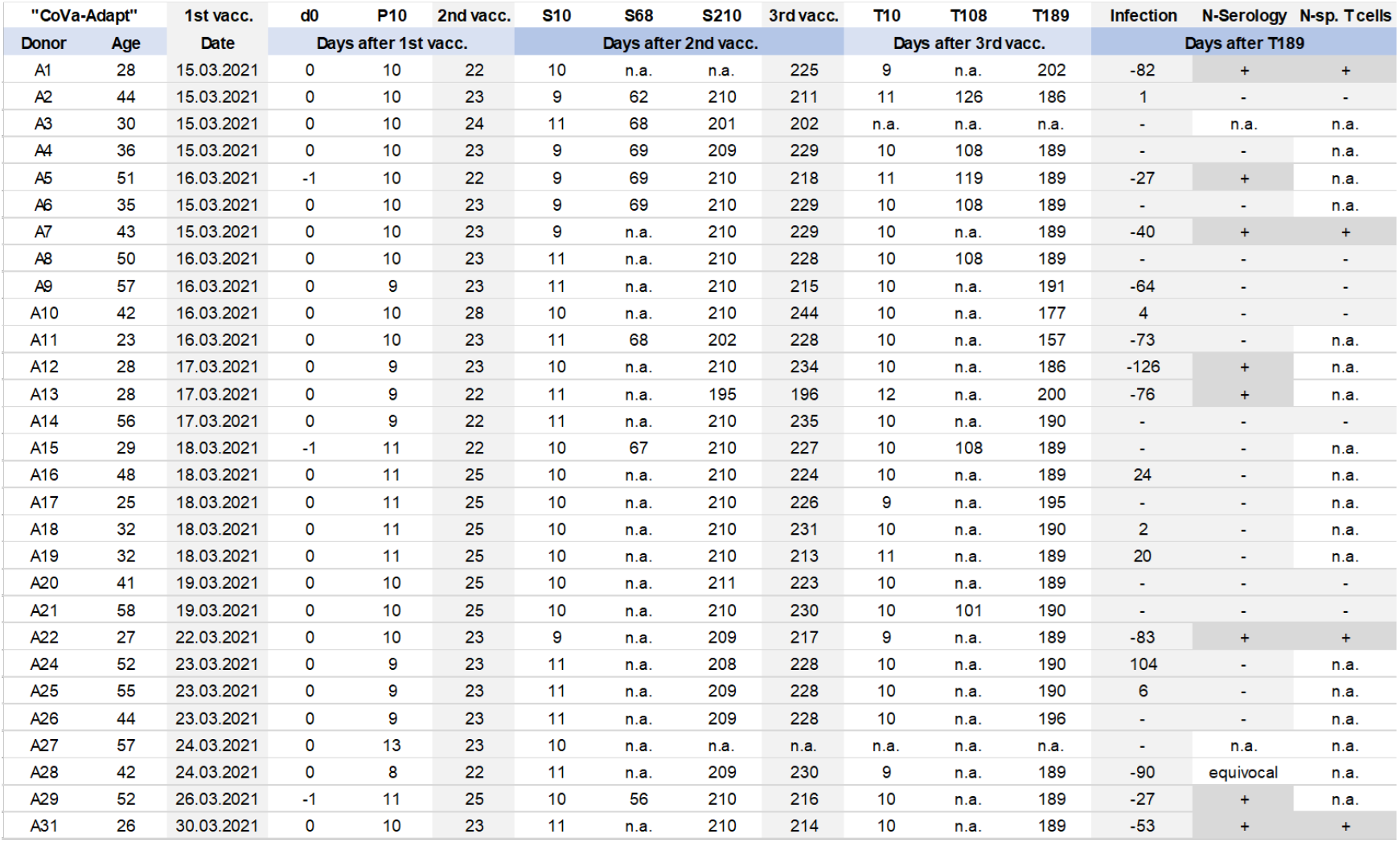
CoVa-Adapt clinical study cohort with sampling time points and information on breakthrough infections. 29 healthy donors, 23-58 years old of European Caucasian ethnicity and normal weight received a three-dose mRNA vaccination regimen with Comirnaty. Two donors (A10 and A28) received mRNA-1273 vaccine from Moderna as a third vaccination. Time intervals between immunizations were 22-28 days between 1^st^ and 2^nd^ dose, and 196-244 days between 2^nd^ and 3^rd^ dose. Blood was sampled before the 1^st^ dose (d0), 8-13 days post 1^st^ dose (P10), 9-11 days post 2^nd^ dose (S10), 195-211 days post 2^nd^ dose (S210), 9-12 days post 3^rd^ dose (T10) and 157-202 days post 3^rd^ dose (T189). Additional blood samples were collected for eight donors 56-69 days after 2^nd^ dose (S68) and for seven donors 101-126 days after 3^rd^ dose (T108). Eleven donors self-reported SARS-CoV-2 breakthrough infections after T10 which were confirmed by PCR tests. Seven additional donors experienced breakthrough infections after T189. For each donor, SARS-CoV-2 nucleocapsid (N) serology and detection of nucleocapsid-specific (N-sp.) T cells by ELISpot is indicated as positive (+) or negative (-), n.a indicates not acquired.

**Fig 1:**
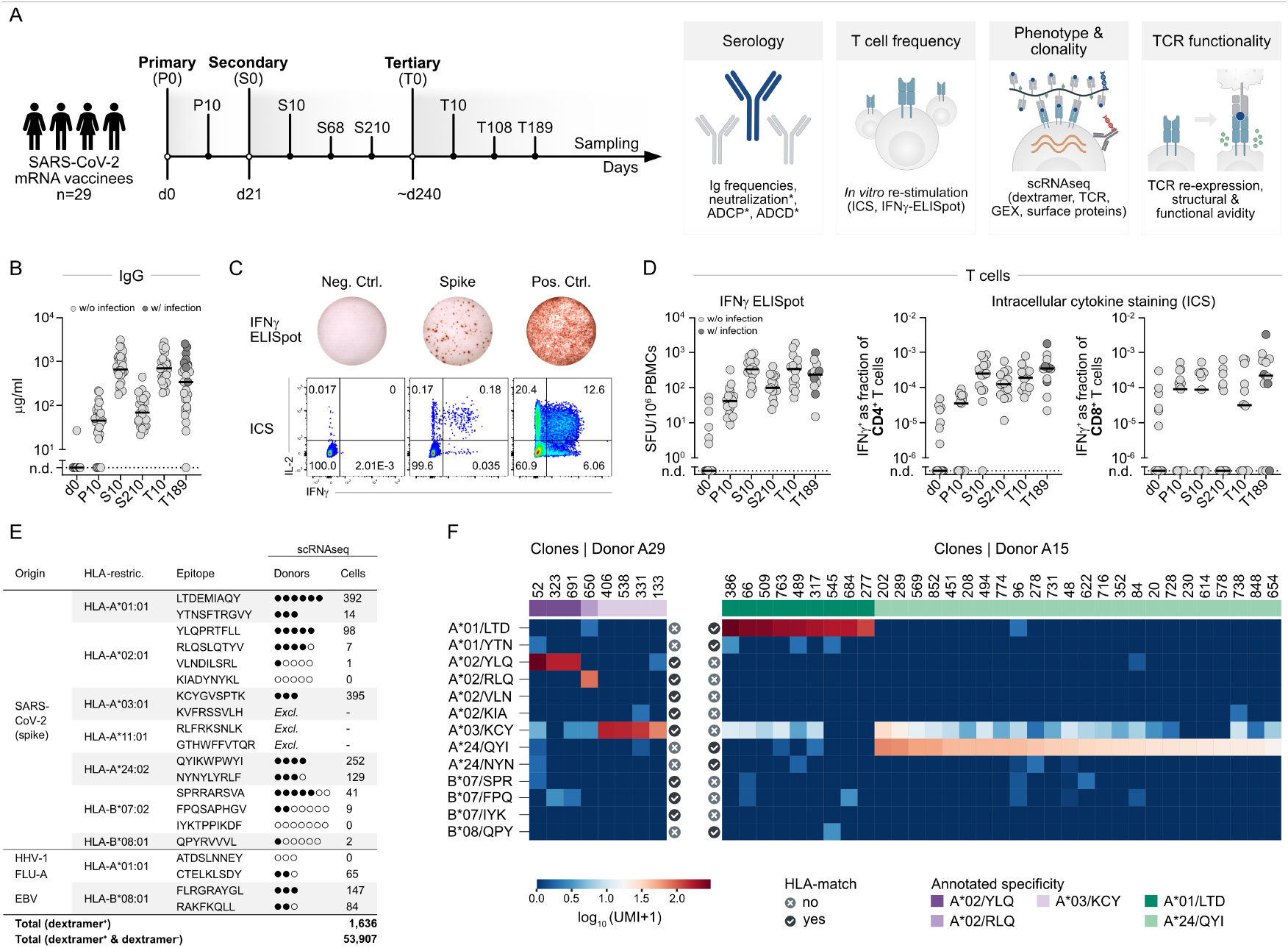
Identification of SARS-CoV-2-specific T cell responses. **A** CoVa-Adapt study design and sample collection scheme. For all donors, PBMCs and serum were collected at day 0 (d0), 10 days after primary (P10), 10 and 210 days after secondary (S10, S210), and 10 and 189 days after tertiary (T10, T189) vaccination. For selected donors, PBMCs were additionally sampled 68 days after secondary (S68, n=8) and 108 days after tertiary (T108, n=7) vaccination. 11 donors experienced a breakthrough infection between T10 and T189 with corresponding positive nucleocapsid serology and are marked in dark grey in sub-figures B-D. Vaccination-induced antibody-and T cell responses were characterized for most donors on a quantitative level. Selected CoVa-Adapt donors (n=13) were subjected to in-depth characterization of CD8 T cell quality using scRNAseq and TCR functional testing. Analyses marked with asterisk are reported in Irrgang et al. **B** Spike-specific IgG in serum quantified by a flow cytometric assay using full-length spike protein as targets. Samples below the lower limit of quantification (15.8 μg/ml) were set to not detected (n.d.). Data points represent individual donors (n=27-29 per time point), solid lines indicate the mean. **C, D** Identification of spike-specific CD4 and CD8 T cells after 20h of *in vitro* re-stimulation of donor PBMCs with 15-mer peptides covering the complete wild type spike protein. Peptides were provided in two sub-pools S1 (Fig. 1C, D) and S2 (Suppl. Fig. 1B); negative control (Neg. Ctrl) = solvent, positive control (Pos. Ctrl) = PMA/ionomycin. Flow cytometric intracellular cytokine staining (ICS) plots are pre-gated on living CD4^+^ lymphocytes (C). Quantification (D) of spot forming units (SFU) for IFNγ ELISpot (left) and IFNγ^+^ T cells as a fraction of living CD4 (middle) or CD8 (right) T cells for ICS, data points represent individual donors (n=12-19 per time point), solid lines indicate the mean. Samples without IFNγ^+^ T cells above the negative control were set to not detected (n.d.). **E** List of HLA-I dextramers presenting SARS-CoV-2 spike, HHV-1, Flu-A, and EBV epitopes used in this study. HLA-matched donors are indicated as filled circles if epitope-specific cells were detected and as unfilled circles if no cells were detected. Number of detected epitope-specific T cells across all HLA-matched donors is indicated. Excl. = excluded dextramers. **F** Representative heatmaps showing average UMI counts of detected clones with assigned epitope-specificity for all stained dextramers in one sequencing experiment for CoVa-Adapt donors A29 and A15. For each epitope, donor-dependent HLA-matching is indicated.

We next characterized the quality of the CD8 T cell response with epitope-specific resolution. To this end, we combined peptide human leukocyte antigen (pHLA) multimers (“dextramers”) presenting one of 16 previously validated spike epitopes^24,31–33,46^ with single-cell RNA sequencing (scRNAseq), single-cell TCR sequencing (scTCRseq), and cellular indexing of transcriptomes and epitopes by sequencing (CITEseq) of 130 surface antigens (**Fig. 1A; Suppl. Table 2**). All epitopes were presented on one of seven HLA alleles that are prevalent in Western and Asian populations. We also added four dextramers presenting epitopes of herpes simplex virus (HHV-1), influenza A virus (Flu-A), and Epstein-Barr virus (EBV) (**Fig. 1E**). All dextramers had an epitope-specific DNA barcode and shared a fluorochrome for fluorescence-activated cell sorting (FACS). Using FACS, epitope-specific CD8 T cells were enriched (not purity-sorted) from PBMCs to ensure maximum yield of epitope-specific cells while simultaneously including epitope-unspecific cells for contextualization (**Suppl. Fig. 2A**). Three scRNAseq experiments were conducted for a total of 14 donors and seven time points after vaccination (**Suppl. Table 2**). Initially included dextramers for epitopes restricted by HLA-A*11:01 (RLF and GTH) and HLA-A*03:01 (KVF) generated an unspecific signal in donors that were HLA-mismatched and were subsequently excluded. For the remaining 13 SARS-CoV-2 spike and four control virus epitopes, we analyzed the number of unique molecular identifiers (UMIs) for each dextramer per TCR clonotype (**Fig. 1F, Suppl. Fig. 2B**). This revealed different signal-to-noise ratios for individual dextramers and experiments. For example, a set of clones with high UMI counts was detected for HLA-A*01:01/LTD and HLA-A*02:01/YLQ only in donors with matching HLA. In contrast, HLA-A*03:01/KCY, showed a broad staining of multiple clones with intermediate UMI counts also in HLA-mismatched donors. However, a group of KCY-specific clones with high UMI counts could still be identified in HLA-matched donors, indicating specific staining above a higher UMI background. For the control virus dextramers HLA-A*01:01/CTE (Flu-A), HLA-B*08:01/FLR (EBV) and HLA-B*08:01/RAK (EBV), elevated UMI counts were detected for several clones (**Suppl. Fig. 2B**). For other spike epitopes and HHV-1, no or only few specific clones could be detected.

To determine optimal UMI cut-offs, we analyzed UMI counts of individual cells belonging to one clonotype for each dextramer (**Suppl. Fig. 2C**). This revealed clone-specific increases in dextramer UMI counts. To refine the annotation of dextramer “positivity”, we implemented additional purity criteria for cells and clones (see Methods; **Suppl. Table 3**). This separated cells into dextramer-positive and dextramer-negative groups with only little or no overlap in UMI distributions for most specificities (**Suppl. Fig. 2D-E**). Interestingly, for the reported^47^ immunodominant epitope HLA-A*24:02/QYI, we could detect many T cells binding dextramers with a seemingly clean signal-to-noise ratio (**Fig. 1F**) in HLA-matched but also mismatched donors (**Suppl. Fig. 2D-E**) which is in line with the recently questioned specificity of this epitope to SARS-CoV-2^38^.

In summary, spike-specific CD8 T cells could be deconvoluted into epitope-specific responses in a multiplexed manner.

### Phenotypes of SARS-CoV-2 epitope-specific CD8 T cell responses

Next, we aimed at obtaining a multi-parametric picture of the phenotypic and clonotypic evolution of epitope-specific CD8 T cell responses. 53,907 cells were recovered post scRNAseq quality control from all donors and across all time points, and were differentiated into 13 phenotypic clusters (**Fig. 2A-B**).

**Fig 2:**
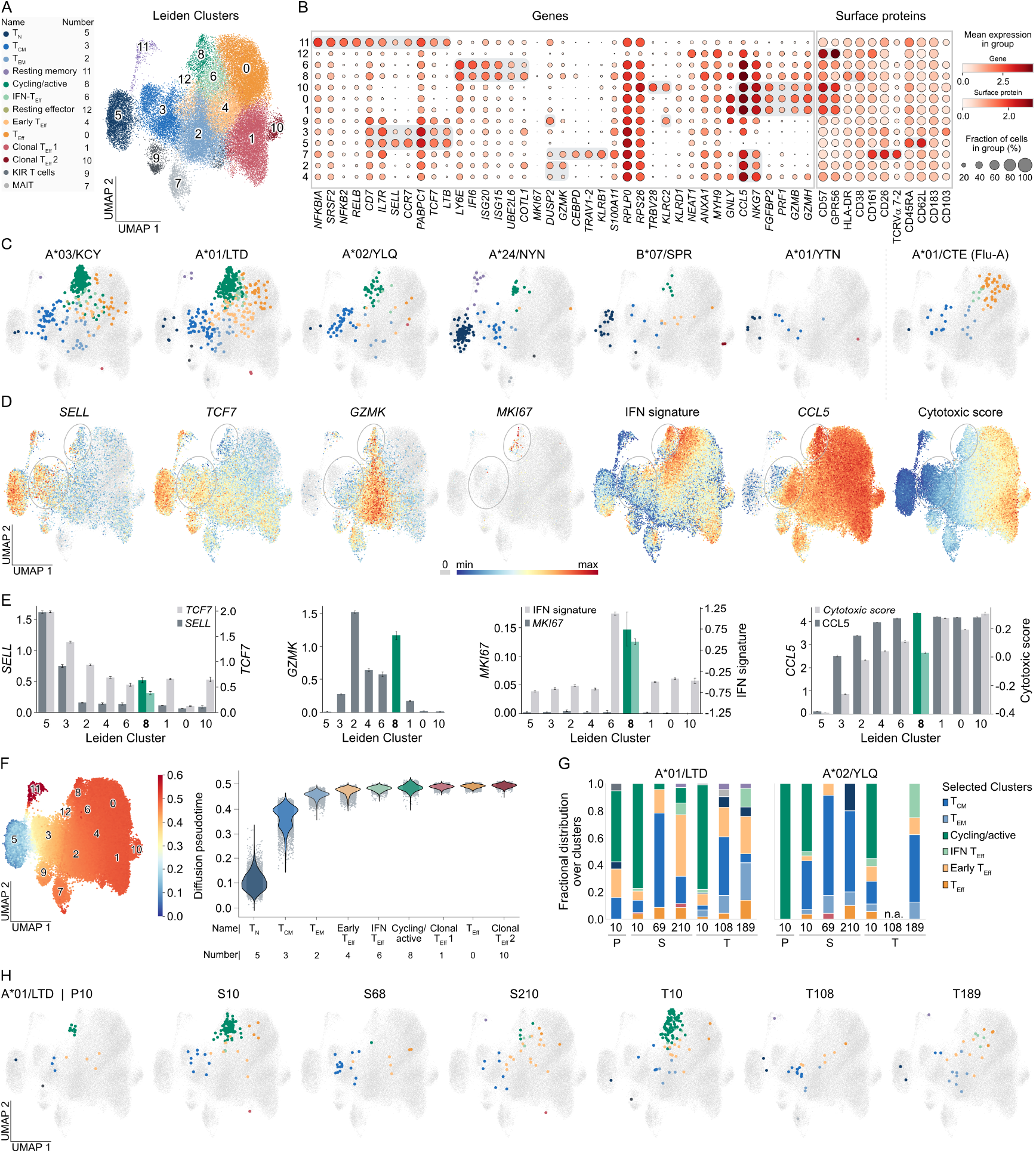
Phenotypic adaptation of epitope-specific CD8 T cell responses throughout repetitive immunization. **A** UMAP with Leiden clusters (cluster numbers in UMAP, cluster names and numbers on the left) of CD8 T cells enriched for pHLA dextramer-binding (Suppl. Fig. 2A) from three independent scRNAseq experiments (n = 53,907 cells). **B** Dot plots of log-normalized expression of representative genes and surface proteins per cluster. Selected genes and proteins are highlighted in grey. Numbers on the left indicate cluster number. **C** Visualization of SARS-CoV-2 spike and Flu-A (A*01/CTE) epitope-specific T cells of all HLA-matched donors (A*03/KCY: n=3, A*01/LTD: n=6, A*02/YLQ: n=5, A*24/NYN: n=4, B*07/SPR: n=7, A*01/YTN: n=3, A*01/CTE: n=3) and screened time points, colored based on their cluster annotation. Cells without the indicated epitope-specificity are shown in grey. **D, E** Log-normalized expression of selected genes and scores. Cells in UMAP are shown in grey for log-normalized gene expression of 0. Cycling/active and T_CM_ clusters are highlighted. (D). For the sake of clarity, quantification was only performed for conventional T cell clusters (i.e., clusters 5, 3, 2, 4, 6, 8, 1, 0, and 10) (E), which are ordered by pseudotime with cycling/active cluster 8 highlighted. Bars with error show the mean and 95% confidence intervals. **F** Visualization of pseudotime on UMAP with indicated cluster numbers (left) and across all conventional T cell clusters with individual cells depicted in grey (right). Starting point for the calculation of pseudotime was in cluster 5. Color gradient was clipped at pseudotime of 0.6 for the sake of clarity (outlier behavior of cluster 11). Clusters with indicated name and number are ordered by pseudotime (right). **G, H** Quantified fractional distribution over all clusters (G) and UMAP visualization of A*01/LTD (H) and A*02/YLQ-specific T cells (Suppl. Fig. 4) of HLA-matched CoVa-Adapt donors (n=5 for A*01/LTD and A*02/YLQ) at individual time points after primary (P), secondary (S) and tertiary (T) vaccination. Colors represent cluster location of epitope-specific cells at the respective time points. Cells without the indicated epitope-specificity ar6e shown in grey. Lack of acquisition of a sample at a certain time point is indicated with not acquired (n.a.).

Phenotypic clusters were annotated using surface protein and gene markers (**Suppl. Table 4**) as naïve T cells (T_N_, cluster 5), central memory T cells (T_CM_, 3), effector memory T cells (T_EM_, 2), cycling/active T cells (8), effector T cells with an interferon sensing signature (IFN T_Eff_, 6), early T_Eff_ (4), T_Eff_ (0), and clonal T_Eff_ (1, 10). Additionally, MAIT (7), suppressive KIR T cells^48^ (9), and two small subsets of resting memory (11) and resting effector (12) T cells were identified.

Combining all time points, T cells specific for LTD, YLQ, and KCY were predominantly found in the cycling/active cluster, but also among T_CM_, IFN T_Eff_ and early T_Eff_ (**Fig. 2C, Suppl. Fig. 3A-B**). For NYN and SPR epitopes, we could detect fewer cells, consistent with a previous report^47^. Remarkably, NYN- and SPR-specific T cells did show some recruitment into the cycling/activated cluster, albeit many cells were also retained in the T_N_ cluster. For seven further epitopes, we detected no or only minute cell numbers, which were then mostly confined to T_N_ or T_CM_ clusters (**Fig. 2C, Suppl. Fig. 3A**). Thus, with decreasing immunogenicity, we observed not only less clonal expansion and differentiation into effector subsets in absolute numbers, but also a higher relative share of early differentiated cells, consistent with previous mouse studies^14,49,50^. Interestingly, QYI-binding cells were found among suppressive CD8 KIR T cells^48^ (**Suppl. Fig. 3A**). While no HHV-1-specific T cells could be detected (data not shown), Flu-A (CTE)-specific cells covered the entire differentiation spectrum, whereas EBV (RAK, FLR)-specific cells were phenotypically synchronized towards T_EM_, IFN T_Eff_ and early T_Eff_ cells (**Fig. 2C, Suppl. Fig. 3A**,**C**).

We used common markers and gene signatures to contextualize cellular differentiation degrees and activation states. Consistent with cellular differentiation status as captured through pseudotime, less differentiated cells expressed *SELL* and *TCF7*, mid-differentiated cells expressed *GZMK* and terminally differentiated cells expressed *CCL5* as well as increasing levels of a CD8 cytotoxic score^51^ (**Fig. 2D-F, Suppl. Fig. 3D**). Furthermore, cells within the cycling/active cluster expressed *MKI67* and high levels of an IFN signature^52^, with the latter extending into the IFN T_Eff_, cluster.

For the T_CM_ cluster, two separate phenotypes were visible throughout the dataset: *SELL*^high^ *CCL5*^low^ less differentiated (top left) and *SELL*^high^ *CCL5*^high^ more differentiated (bottom right) T_CM_ cells (**Fig. 2A,D**). T cells specific for immunodominant epitopes LTD, YLQ, and KCY showed enrichment of cells at the intersection of these two sub-clusters of T_CM_ cells (**Fig. 2C**). Cycling/active cells expressed the highest levels of HLA-DR and CD38 surface protein, confirming their activated phenotype (**Fig. 2B**). Within this cycling/active cluster, we observed graded and opposing expression patterns of the IFN signature and *SELL* (**Fig. 2D**), indicating that cycling/active cells encompass various activation and differentiation states.

The highly standardized character of our CoVa-Adapt cohort enabled us to zoom in on defined time points after vaccination and characterize the phenotype and clonotype of epitope-specific T cells longitudinally. For the immunodominant epitopes LTD, YLQ, and KCY, only few cells were detected at the time point P10, which were phenotypically dominated by cycling/activated cells (**Fig. 2G-H, Suppl. Fig. 4**). At S10, cell numbers of epitope-specific T cells increased in line with previous quantitative characterizations (**Fig. 1D, Suppl. Fig. 1 B-C**). Phenotypically, we again detected predominantly cycling/active cells, while a smaller but distinct subset of less differentiated T_CM_ cells was also retained. Several months after the 2^nd^ vaccination at the early (S69) and late (S210) memory time points, almost none of the spike epitope-specific T cells were active or cycling anymore, whereas less differentiated T_CM_ cells prevailed. Notably, at the late memory time point S210, epitope-specific T cells also adapted a more terminally differentiated phenotype (early T_Eff_ and IFN T_Eff_).

After the 3^rd^ vaccination, epitope-specific cells underwent a similar, but not identical, phenotypic evolution. At late memory (T189), a higher fraction of LTD-specific T_EM_ cells emerged (**Fig. 2G-H**). Also, across the acute time points (P10, S10, T10), epitope-specific cells shifted within the cycling/active cluster with each vaccination, according to the previously described differentiation and activation gradient (**Fig. 2D**). The IFN response signature followed the same transcriptional shift (**Suppl. Fig. 5A-B**). This is consistent with an important role of type I interferons for CD8 T cell responses following mRNA vaccination^53^, and suggests the usefulness of IFN response signatures^51^ as “molecular clocks” in vaccination-induced CD8 T cells.

Cells specific for less immunodominant SARS-CoV-2 epitopes were irregularly detected at individual time points, and mostly maintained their less differentiated phenotype (**Suppl. Fig. 4**). Overall, phenotypic kinetics were consistent with changes in pseudotime over real time after vaccination (**Suppl. Fig. 5C**) and reproducible across different individual donors (**Suppl. Fig. 6A-C**).

Finally, we compared the phenotypic results from our CoVa-Adapt donors to a hypervaccinated individual from Magdeburg (HIM)^54^. At day 189 after 215 SARS-CoV-2 vaccinations (X189), HIM’s LTD-specific T cells were phenotypically skewed towards terminal T_Eff_ cells compared to regular responses 189 days after three vaccinations, with similar fractions of early and IFN T_Eff_ cells and a lower, albeit preserved, fraction of less-differentiated T cells (**Suppl. Fig. 6D**).

Taken together, CD8 T cells recognizing immunodominant spike epitopes were more differentiated and active in the acute phase after each vaccination, while less differentiated subsets with stem-like potential were preserved throughout the immune response.

### Epitope-specific TCR repertoires are polyclonal after SARS-CoV-2 vaccination

To investigate cellular qualities conveyed by the antigen receptor, we explored changes in the TCR repertoire. Clonal expansion generally increased with phenotypic differentiation, which was also reflected by an increasing clonality (Gini index) of the TCR repertoire (**Fig. 3A-B**). Within the cycling/active cluster, the largest clone sizes were observed at the top of this cluster (**Fig. 3A**), in line with the previously described differentiation and activation gradient (**Fig. 2D**). Clonal expansion and clonality within this cluster were higher than for T_CM_, but lower compared to T_EM_ cells.

**Fig 3:**
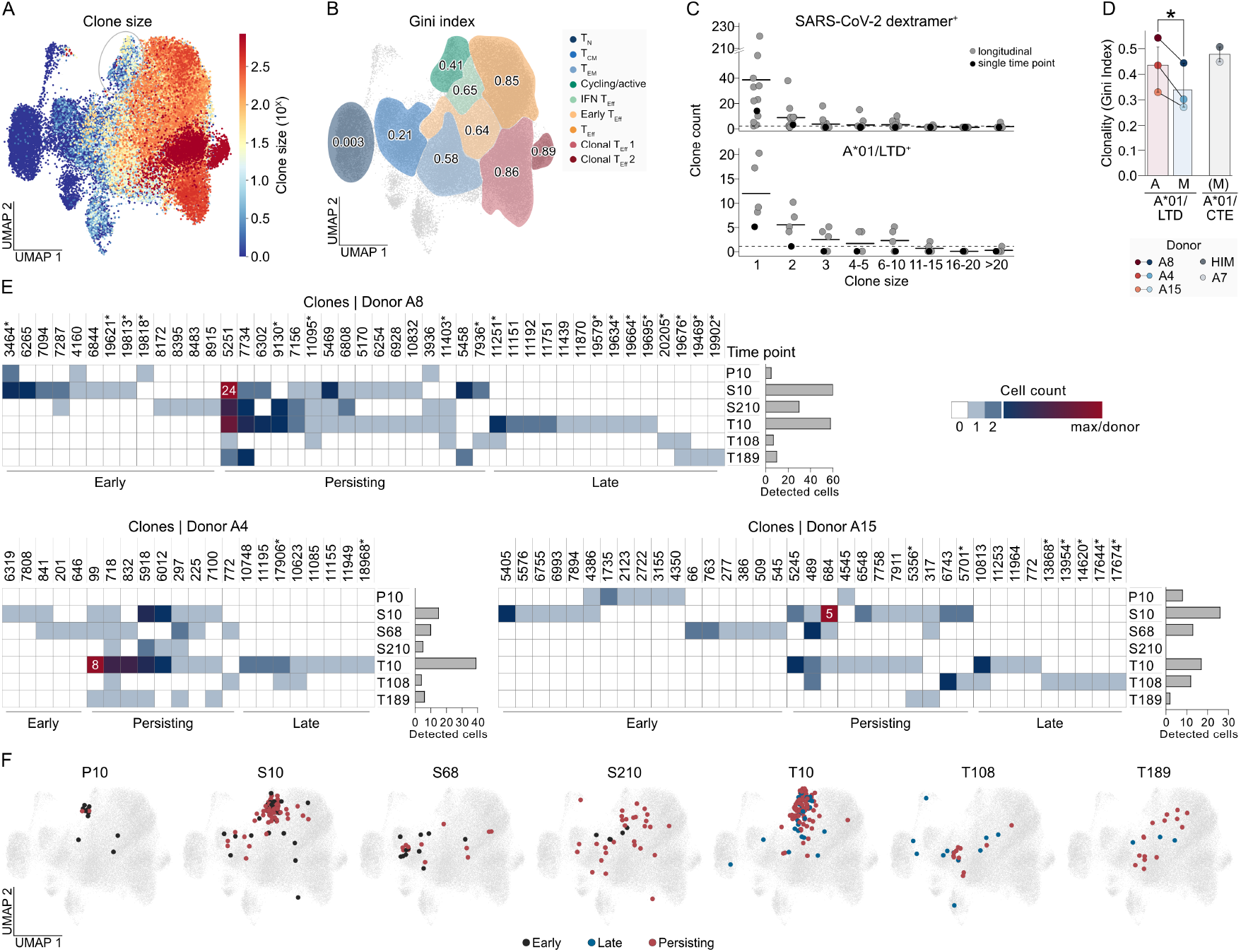
Polyclonality of epitope-specific TCR repertoires. **A** Visualization of clone size for all cells on UMAP. Cycling/active cluster is highlighted. **B** Gini index (high values indicate high clonality, low values indicate high evenness) was calculated over all clones of selected clusters. **C** Number of all SARS-CoV-2 spike dextramer^+^ (top) and A*01/LTD dextramer^+^ clones (bottom) of HLA-matched donors with the indicated clone sizes. Data points represent individual donors and solid lines indicate the mean. Dotted line: clone count = 1. Donor A7, for which only T189 was screened, is indicated in black. **D** Gini index of A*01/LTD dextramer^+^ cells detected in the acute (A: S10/T10) and memory (M: S68/S210/T108/T189) phase. Gini index for A*01/CTE (Flu-A) dextramer^+^ cells from two additional donors is shown for comparison; “(M)” indicates presumed memory phase as no recent influenza infection was reported. Data points represent individual donors, bars with error bars show the mean and 95% confidence interval. Statistical testing by paired t-test. * p<0.05. **E** Detected A*01/LTD dextramer^+^ clones across all scRNAseq experiments are shown for three HLA-matched donors. Color gradient indicates cell counts at individual time points. The maximum detected cell number per donor is indicated in the heatmap. Clones are ordered by detection pattern indicated at the bottom and by cell count at T10. Bar graphs (on the right of each heatmap) show total number of detected A*01/LTD dextramer^+^ cells per time point and donor. Clones marked with asterisk were only detected by re-sequencing and not functionally characterized. **F** Visualization of detected A*01/LTD dextramer^+^ cells from donors A4, A8, and A15 on UMAP at individual time points after each vaccination. Cells are colored based on detection pattern.

We next investigated T cell responses to the immunodominant epitope LTD in individual donors. In total, we detected 121 unique LTD-specific clonotypes from four longitudinally screened donors and one single-time point donor. As for all spike epitopes, the majority of LTD-specific clones were singlets, characterizing the repertoires as diverse even after repetitive vaccination (**Fig. 3C**). Only few clones dominated LTD-specific repertoires in individual donors with cell numbers exceeding five cells, mirroring repertoire dynamics described in mouse models of infection^14^. Despite this overall high diversity, we observed clonal skewing in the acute phase after each vaccination, which decreased again in the memory phases (**Fig. 3D**).

Among the longitudinally screened donors, donor A16 harbored an unusually small repertoire. We therefore focused on LTD-specific repertoires from donors A4, A8, and A15 to evaluate the evolution of epitope-specific TCR repertoires after vaccination (**Fig. 3E**). Samples were, if possible, re-sequenced in multiple experiments to increase and solidify cell numbers and detection dynamics of individual clones. As observed before, the number of detected LTD-specific cells peaked at acute time points after 2^nd^ and 3^rd^ vaccination (S10, T10) and decreased at early (S68, T108) and late memory (S210, T189) time points. Across the three LTD repertoires and also for YLQ-specific cells in donor A29 (**Suppl. Fig. 7A**), clones could be divided into three detection patterns: early, late and persisting – depending on their detection before, after, or before and after 3^rd^ vaccination. In line with previous reports^55^, most persisting clones did not appear before the 2^nd^ vaccination (but only from S10 onwards) and were more expanded especially after the 3^rd^ vaccination (T10) (**Fig. 3E**). While some clones were expanded and appeared at multiple time points, each antigen encounter resulted in the detection of many additional, unique TCR clonotypes throughout the repertoires (**Suppl. Fig. 7B**). Newly emerging (“late”) clones contributed to the maintenance of a high TCR repertoire diversity even after three vaccinations (**Fig. 3E**). Notably, for persisting TCRs with substantial expansion at T10 like clone 5251 (donor A8), 99 (donor A4), or 489 (donor A15), we could observe the same phenotypic dynamics that we previously described for epitope-specific populations as a whole (**Suppl. Fig. 7C**). This indicated that individual clonotypes did not occupy specific phenotypic niches. Also, we could not detect phenotypic differences for clones with early, late or persisting detection patterns (**Fig. 3F**).

After hypervaccination, LTD-specific clones showed a higher degree of clonal expansion^54^. However, this repertoire still consisted of multiple unique clones and therefore also maintained clonal diversity (**Suppl. Fig. 7D-E**). In summary, epitope-specific TCR repertoires underwent dynamic changes after each vaccination, with longitudinally dominating clones being mostly formed after the 2^nd^ vaccination. Through persistence and new emergence of clonotypes, repertoire diversity was maintained throughout repetitive immunization.

### Recruitment of high-functionality CD8 T cell clones after SARS-CoV-2 vaccination

Having documented the precise clonal dynamics of epitope-specific TCR repertoires, we were in a position to address the fundamental question whether and how TCR avidity determined clonal recruitment and maintenance of CD8 T cells in the human system. To this end, we re-expressed 106 TCRs covering four LTD- and one YLQ-specific repertoires and characterized their avidity landscape (**Suppl. Tables 5-6**). The investigated clones were detected after the first two of three sequencing experiments. Clones that only emerged upon re-sequencing were not included, but did not represent expanded or longitudinally re-occurring clones (marked by asterisks in **Fig. 3E**).

To analyze TCR functionality under near-physiological conditions, we introduced each TCR into primary human T cells via CRISPR/Cas9-mediated orthotopic TCR replacement (OTR) (**Suppl. Fig. 8A**)^56–59^. This technique allows the TCR to be precisely integrated into the endogenous TCR gene locus, preventing overexpression and enabling physiological regulation of the transgene. We were able to re-express 104/106 TCRs with sufficient knock-in efficiency (protein expression; **Suppl. Fig. 8B-D**). Each TCR was characterized for its ability to bind fluorophore-conjugated pHLA multimers (which entailed streptavidin as a backbone and no DNA barcode) and its sensitivity to exert effector functions (IFNγ upregulation, TCR downregulation) after co-culture with peptide-loaded antigen-presenting cells (APCs). “Functional avidity” was then documented as the peptide concentration at which half-maximum effector function was reached (“EC_50_ value”; **Fig. 4A-C, Suppl. Fig. 9**).

**Fig 4:**
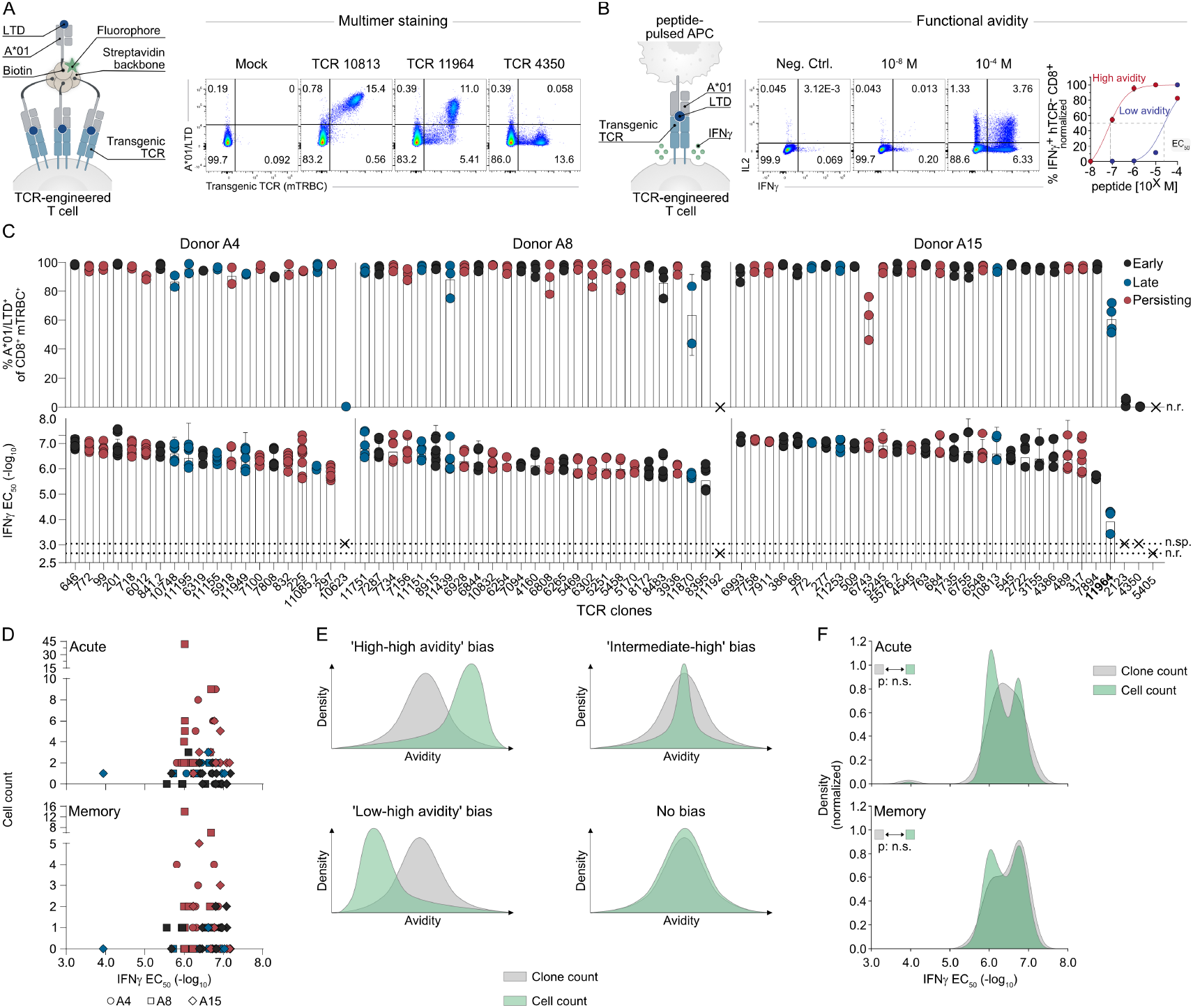
High functionality of epitope-specific T cell receptors irrespective of clonal expansion. **A** A*01/LTD dextramer^+^ TCRs were transgenically re-expressed in primary human T cells by CRISPR/Cas9-mediated orthotopic TCR replacement (OTR) with a murine constant region (mTRBC) to be distinguishable from the endogenous TCR (hTCR) and stained with A*01/LTD multimers on day 11 after OTR. Primary data for representative TCR clones are pre-gated on living CD8 lymphocytes. Mock: unedited T cells. **B** TCR-engineered T cells were co-cultured with APCs loaded with LTD peptide ranging from 10^-8^ to 10^-4^ M in a ratio of 1:1 for 4 h. Primary data are pre-gated on living CD8 hTCR^-^ lymphocytes. Representative dose-dependent stimulation is shown for clone 10813 (red) and 11964 (blue) with quantification of peptide sensitivity by EC_50_ values. Negative control (Neg. Ctrl) = solvent. **C** Quantification of A*01/LTD multimer^+^ cells of living CD8 hTCR^-^ mTRBC^+^ lymphocytes for donor A4 (n=2-4 technical replicates from 2-4 experiments per TCR), A8 (n=2-3 from 2-3 experiments per TCR), and A15 (n=2-4 from 2-4 experiments per TCR) (top). Quantification of peptide sensitivity (EC_50_) determined by dose-dependent IFNγ upregulation of living CD8 hTCR^-^ lymphocytes for donor A4 (n=4-10 from 2-5 experiments per TCR), A8 (n=4-6 from 2-3 experiments), and A15 (n=4-6 from 2-3 experiments) (bottom). TCRs are ordered by increasing EC_50_ value (decreasing functionality) from left to right and colored based on their detection pattern. Numbers at the bottom indicate TCR identifiers, TCRs for which a second α-or β-chain was identified by scRNAseq and functionally tested are identified with “.2” at the end. Data points represent technical replicates, bars with error bars show the mean +/-s.d.. TCRs that were not re-expressed by OTR or did not show IFNγ upregulation above the negative control at the highest peptide concentration were labeled as not re-expressed (n.r.) or not specific (n.sp.), respectively. TCR 11964 with ultra-low avidity (EC_50_ of -log_10_ 3.94 M) is highlighted in bold (4^th^ TCR from right). **D** Correlation of EC_50_ values with cell counts of respective clones in the acute (P10/S10/T10) and memory (S68/S210/T108/T189) phase after vaccination. Data points represent individual TCRs, symbols represent individual donors, and color represents detection pattern of each TCR clone. **E** Theoretical models how a cell count distribution is skewed towards expansion of ‘high-high avidity’ (EC_50_ of around -log_10_ 7 M), ‘intermediate-high avidity’ (EC_50_ -log_10_ 6.5 M), or ‘low-high avidity’ (EC_50_ -log_10_ 6 M) TCR clones, or if TCR avidity does not bias clonal expansion (‘No bias’). Density on the y-axis indicates clone count for clone count distribution (grey), or cell count per clone for cell count distribution (green). **F** Clone count (grey) and cell count (green) distribution for A*01/LTD-specific clones fr1om0 repertoires of donor A4, A8, and A15. Clone and cell count is depicted for acute (P10/S10/T10) and memory (S68/S210/T108/T189) phase after vaccination. Statistical comparison of clone and cell count distribution was performed by Kolmogorow-Smirnoff test, n.s., not significant.

These analyses confirmed 98/106 (92%) of TCRs to be epitope-specific and reactive, demonstrating the accuracy of our workflow to identify epitope-specific CD8 T cells through scRNAseq with DNA-barcoded dextramers. CD8 T cells with the same epitope specificity showed co-clustering in TCR sequence distance (**Suppl. Fig. 10A**). This enabled us to identify most functional alpha (10/13) or beta chains (8/8) at first guess when epitope-specific T cell clonotypes showed two alpha or beta chains^60^. Of note, we could not validate the QYI specificity for three QYI dextramer-binding TCRs, confirming the suspected unspecific signal for this epitope (**Suppl. Fig. 8E, Suppl. Fig. 11**).

Overall, all three LTD repertoires consisted of highly functional T cell clones (EC_50_ mostly > -log_10_ 6 molar (M); note that high avidities are indicated by high -log_10_ EC_50_ values) with minimal avidity differences between TCRs (**Fig. 4C, Suppl. Fig. 9A**). Only one clone, TCR 11964 (donor A15; marked in bold), showed a markedly lower functional avidity (EC_50_ of -log_10_ 3.94 M) and reduced pHLA multimer binding. Similar findings were obtained in a YLQ-specific repertoire, which was also characterized by highly functional T cell clones with avidities that were generally slightly higher than LTD avidities, at a level comparable to a cytomegalovirus-specific control TCR^58^. Again, only one clonotype (8191) showed a markedly reduced TCR avidity (**Suppl. Fig. 9B**).

To investigate whether subtle differences in functionality among generally high avidities have biological effects, we categorized avidity levels as “low-high”, “intermediate-high” and “high-high” with EC_50_ of around -log_10_ 6, 6.5, or 7 M, respectively. In all characterized repertoires, persisting and longitudinally dominating clones were not enriched for “high-high” avidities, but rather distributed across the entire spectrum of overall high TCR functionalities (**Fig. 4C, Suppl. Fig. 9A-B**). Remarkably, TCR avidity was also not correlated to clonal expansion at acute or memory time points (**Fig. 4D**). Instead, some clones that were particularly expanded possessed avidities in an “intermediate-high” range (EC_50_ of around -log_10_ 6.5 M). However, other clones with such avidities were not clonally expanded. Even after 215 vaccinations and a more pronounced clonal skewing (donor HIM, **Suppl. Fig. 7D-E**), there were no remarkable avidity differences in the recruited TCR repertoire that could be linked to differential clonal expansion (**Suppl. Fig. 9C**). Furthermore, neither detection pattern, clonal expansion nor avidity were associated with TCR sequence similarity clusters (**Suppl. Fig. 10B-E**). While IFNγ release and TCR downregulation significantly correlated with each other (**Suppl. Fig. 12A**), no or only a mild correlation (depending on the donor) was observed between functional avidity and pHLA multimer binding intensity, which is often used as a surrogate readout for TCR avidity (**Suppl. Fig. 12B**).

Initially, we were surprised that persistent and expanded clones did not show a bias for “high-high” TCR avidities (EC_50_ of around -log_10_ 7 M). To explore whether and how TCR avidity affected clonal expansion, we devised multiple models (**Fig. 4E**). Most clones that were recruited into epitope-specific repertoires showed an “intermediate-high” TCR avidity, whereas fewer clones possessed “low-high” or high-high” avidities. Clone counts thus followed a normal distribution. Next, we figured that a “high-avidity bias” would lead to disproportionate clonal expansion of clones with high TCR avidities, resulting in a skewing of cell count distributions towards higher avidities. Alternatively, “intermediate-high” TCR avidities may drive clonal repertoire evolution. Then, such avidities would disproportionately lead to high clonal expansion, resulting in a narrowing of cell count distributions in the middle (“intermediate-avidity bias”). Finally, clonal expansion of TCR repertoires that generally contain TCRs of high functionalities may also be probabilistic. In this case, the cell count distribution would be proportionate to the clone count distribution, i.e., how many clones are present in the recruited repertoire (“no bias”).

In the acute (P10, S10, T10) and memory (S68, S210, T108, T189) phase after vaccination, clone and cell count distributions of LTD-(**Fig. 4F**) and YLQ-specific (**Suppl. Fig. 13A**) repertoires were highly overlapping. Despite the occurrence of local peaks due to expansion of individual clones, clone and cell count distributions did not differ statistically significantly from each other. The same observation was made for LTD-specific repertoires after hypervaccination and thus 215 potential rounds of clonal selection (**Suppl. Fig. 13B**). These findings are best compatible with a model in which clonal outgrowth is probabilistic and can occur across a wide spectrum of avidities as long as a minimum threshold is surpassed (EC_50_ of around -log_10_ 6 M). Accordingly, the two clones that possessed low avidities (EC_50_ values < -log_10_ 5 M) were not clonally expanded at any time point (clones 11964 and 8191 specific for LTD and YLQ; **Fig. 4D, Suppl. Fig. 13A**). Despite the minimal influence of TCR avidity on clonal expansion within recruited repertoires, a minor avidity increase for LTD-specific populations as a whole was observed from 2^nd^ to 3^rd^ vaccination (**Suppl. Fig. 13C**). This indicates continuous, but moderate maturation of the quality of TCR repertoires on the population-level.

In summary, SARS-CoV-2 vaccination led to the recruitment of highly functional CD8 T cell clones. Only 2/98 LTD-or YLQ-reactive clones harbored low avidities. Within the recruited repertoires, clonal expansion was not tuned by subtle differences in avidity, but rather followed a probabilistic pattern. Clones with “intermediate-high” TCR avidities were thereby most likely to undergo clonal expansion since they represent the majority of clones that are recruited into antigen-experienced repertoires in the first place.

### Active recruitment from the naïve repertoire is governed by TCR avidity thresholds

Characterization of peptide sensitivity for 98 TCR clonotypes with defined epitope specificities provided us with the opportunity to characterize the phenotypic traits of CD8 T cell clones with different avidities at multiple time points after vaccination. Highly functional LTD- and YLQ-specific clones (EC_50_ of > -log_10_ 6 M) efficiently gave rise to differentiated T cells, with no single cell of these clones remaining in the T_N_ cluster after vaccination (**Fig. 5A, Suppl. Fig. 14**). Conversely, two clones (one LTD-specific and one YLQ-specific clone) with markedly lower avidity exclusively maintained a naïve phenotype. Together with a lack of expansion, this indicated that these clones did not enter functional immune responses.

**Fig 5:**
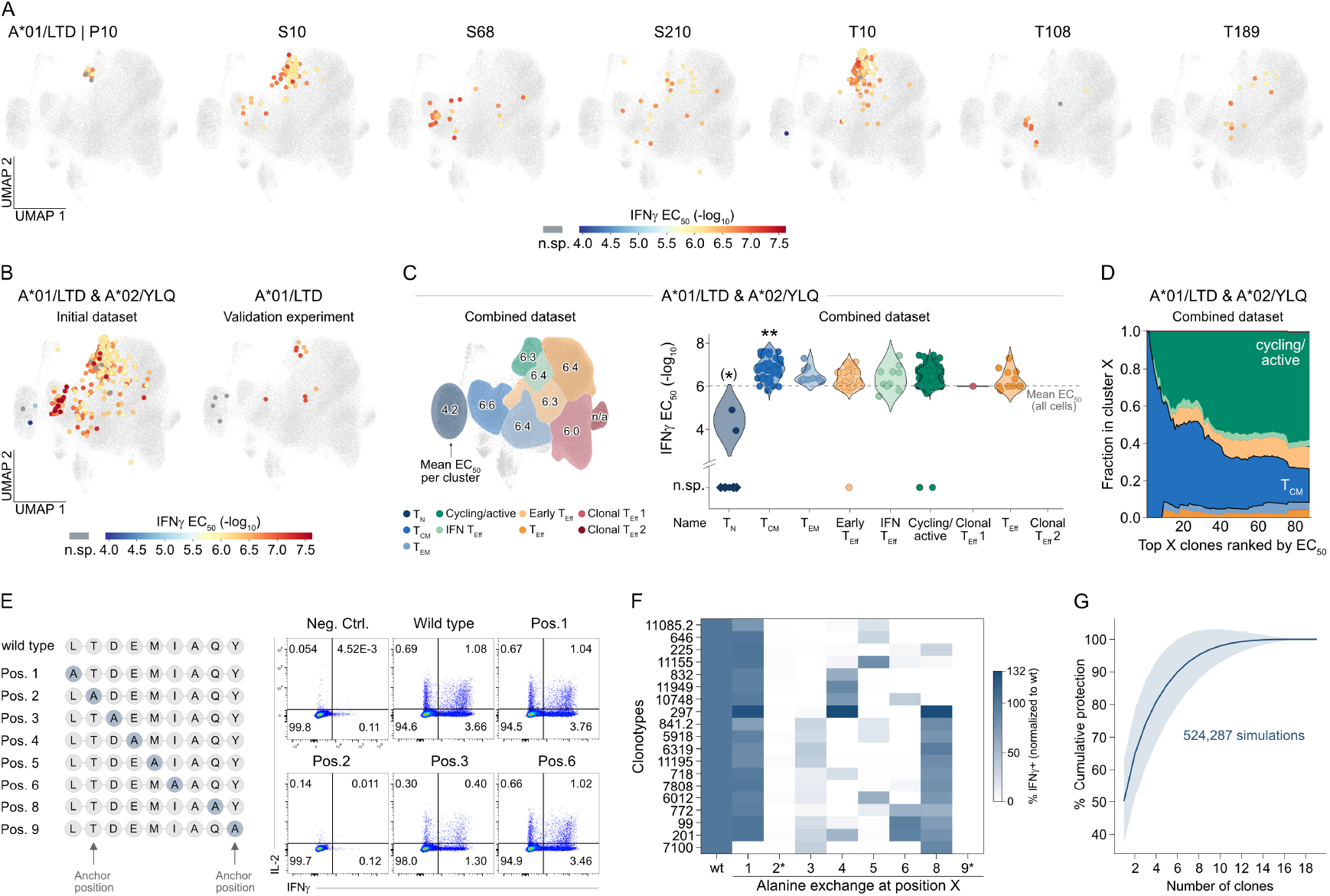
Governance of clonal recruitment from the naïve repertoire by TCR avidity thresholds resulting in polyclonal cross-protective repertoires. **A** Visualization of all functionally characterized A*01/LTD dextramer^+^ TCRs (donor A4, A8, A15) on UMAP at individual time points after each vaccination. Color gradient indicates EC_50_ values. Cells of clones that did not show IFNγ upregulation above the negative control at the highest peptide concentration were labeled as not specific (n.sp.) and are shown in dark grey and circled large dots. Not functionally characterized cells are shown in light grey, uncircled small dots. **B** Visualization of all initially re-expressed and functionally characterized A*01/LTD (donor A4, A8, A15) and A*02/YLQ (donor A29) dextramer^+^ TCRs for all time points pooled (left). Visualization of six additionally characterized A*01/LTD dextramer^+^ TCR clones (right). **C** Mean EC_50_ values over all functionally characterized A*01/LTD and A*02/YLQ dextramer^+^ TCRs for individual clusters on UMAP (left). Colors indicate cluster annotation. n/a indicates not applicable. EC_50_ values for individual A*01/LTD and A*02/YLQ dextramer^+^ cells are shown for conventional T cell clusters (right). Cluster annotation is shown at the bottom. Clusters are ordered by pseudotime. Mean EC_50_ values across all cells are indicated by a dotted line. Statistical testing of EC_50_ per cluster vs. other clusters by Mann-Whitney U test. ** p<0.01, (*) p<0.05 statistically different but in brackets due to few data points. Symbols distinguish cells from the initial (circles) and validation (diamonds) data sets. **D** Fractional distribution across all conventional T cell clusters shown in (C) for cells of top X clones ranked by EC_50_ values (starting with highest functionality TCR on the left). **E-G** Engineered T cells expressing A*01/LTD-specific TCRs from donor A4 were co-cultured with APCs loaded with 10^-5^ M of wild type (wt) LTD or altered peptide ligands (APLs) in a ratio of 1:1 for 4 h. Scheme of APLs (left) ordered by mutated position (alanine highlighted). Representative flow cytometry plots are shown for TCR 99 and pre-gated on living CD8 hTCR^-^ lymphocytes (E). Quantification of IFNγ upregulation normalized to stimulation with wt LTD for each individual TCR. Clones are ordered by reactivity pattern. Anchor positions are indicated with asterisk (F). Degree of protection (y-axis, meaning for how many APLs reactivity is provided excluding mutations at anchor positions) was calculated for different numbers (x-axis) and combinations (data distribution) of TCR clones. TCRs were randomly drawn (524,287 simulations) from the repertoire and the protection score was calculated for each TCR combination for the given number of clones. The reactivity of clone 297 against wt LTD was set as a threshold for a protective reactivity against APLs. The line represents the mean protection score with the area indicating s.d. (G).

To further investigate avidity-dependent clonal recruitment, we re-expressed four additional LTD-specific clones that were detected exclusively in the T_N_ cluster after vaccination and consisted of a single cell each (**Suppl. Fig. 15A**). As controls, two clones (9130 and 11251) with prominent recruitment of cells into the cycling/active cluster were included in this validation experiment. All six clones had not been previously functionally characterized as they were identified in different donors than studied before or only emerged after re-sequencing of samples. Although all six clones were successfully re-expressed, only the TCRs belonging to the two clones with cycling/active cells were able to bind pHLA multimers and upregulated IFNγ after co-culture with LTD-pulsed APCs (**Fig. 5B, Suppl. Fig. 15B-D**). The four other TCRs did not show epitope reactivity, and were thus either falsely dextramer-positive, or of no sufficient avidity to be recruited. This strengthened the hypothesis that epitope-specific T cells with a differentiated (non-naïve) phenotype harbored highly functional TCRs. In contrast, quantification of TCR avidity per phenotypic cluster confirmed that a naïve phenotype was tightly linked to low or no TCR functionality (**Fig. 5C, Suppl. Fig 15E**). This implied that phenotypically naïve T cells that are induced by SARS-CoV-2 mRNA vaccination were indeed antigen-inexperienced or insufficiently primed, unlike, for example, T cells induced by yellow fever vaccination, which is known to induce an antigen-experienced yet naïve-like phenotype^61,62^. Interestingly, spike epitope-specific T_CM_ cells were enriched for a group of cells recognizing their respective epitope with particularly high avidity (**Fig. 5C-D**). This is in line with the concept of preferential preservation of highly functional T cells with stem-like potential^63^. Notably, the differentiation degree of epitope-specific T cells was not linked to TCR similarity clusters (**Suppl. Fig. 10F**).

Active recruitment of epitope-specific CD8 T cell clones into the immune response after vaccination therefore required a threshold level of TCR functionality (EC_50_ of > -log_10_ 6 M). Furthermore, epitope-specific T cells with a particularly high TCR avidity were maintained in stem-like T cell subsets.

### Polyclonal repertoires provide flexibility against mutational escape mechanisms

Across multiple TCR repertoires, we observed a substantial degree of TCR diversity that was maintained after each vaccination. We wondered whether this polyclonality could provide cross-protection against escape mutations. We could not identify mutations of the LTD epitope in naturally occurring SARS-CoV-2 variants, but performed an alanine scan of the LTD peptide to mimic mutational escape. All altered peptide ligands (APLs) were predicted binders for HLA-A*01:01 by netMHCpan-4.1^64^ and used for *in vitro* re-stimulation. The complete LTD-specific repertoire of donor A4, consisting of 19 TCRs, was tested for reactivity against the wild type (wt) epitope and all eight possible APLs (**Fig. 5E-F, Suppl. Fig. 16**). None of the TCRs showed reactivity for APLs mutated at positions 2 and 9, in line with these positions harboring conserved amino acids for HLA-A*01:01-specific peptide presentation^65^. Most, but not all, TCRs were reactive against APLs with mutations at positions 1 or 8 to a similar extent as towards the wt peptide. Interestingly, clone 297, which had the lowest avidity against the wt epitope (**Fig. 4C**), recognized APLs mutated at positions 1, 4 and 8 with increased reactivity (**Fig. 5F, Suppl. Fig. 16**). Overall, all TCRs provided reactivity against at least two APLs while no TCR was able to cover the complete mutational landscape (**Fig. 5F**), demonstrating the necessity of a polyclonal repertoire to counteract unpredictable mutational escape in a flexible manner.

Finally, we investigated how many different TCR clones were required for robust protection. For this, we randomly sampled TCRs from the LTD-specific repertoire and calculated the “cumulative protection” conveyed by increasing repertoire sizes (meaning for how many APLs reactivity is provided, excluding mutations at anchor positions). Clone 297 showed the lowest avidity (**Fig. 4C**) that still led to active recruitment after vaccination (**Fig. 3E**). We therefore set the reactivity of this clone against the wt peptide as a threshold for protective reactivity. Simulations of uniquely combined repertoires revealed that a minimum of 12 clones was required to provide an average cumulative protection of 99.3% (**Fig. 5G**). Of note, 6/7 LTD-or YLQ-specific repertoires for which we obtained longitudinal data entailed at least 13 different clones, suggesting that these immunogenic epitopes induced a degree of polyclonality that is expected to lead to robust cross-protection.

## Discussion

We here describe the longitudinal characteristics of human antigen-specific T cell responses following SARS-CoV-2 mRNA vaccination with single-cell and single-clonotype resolution. After broadly investigating T cell responses, we focused on individual SARS-CoV-2 HLA-I-restricted epitopes for in-depth analysis. Out of 16 epitopes that had been previously reported to be immunodominant and restricted by common HLA alleles, we confirmed strong T cell responses for LTD, YLQ, and KCY. T cell responses against these epitopes showed similar kinetics with regard to their phenotypic evolution, including a prominent recruitment into cycling/active subpopulations shortly after each immunization. During memory time points, antigen-specific T cells encompassed a higher relative fraction of T_CM_ cells with an early differentiation profile, but also maintained a substantial number of more differentiated T_EM_ and T_Eff_ cells, resembling responses induced by influenza virus exposure.

Many studies have investigated the differentiation kinetics of antigen-specific T cells in mouse models and reported similar phenotypes. By single-cell transfers, genetic fate-mapping, and adoptive transfers, it could be shown that early differentiated cells most likely represent *bona fide* memory cells^66^ that give rise to subsequent recall responses^67^. At the same time, terminally differentiated subsets do not suddenly vanish after the antigen is gone^38^, but contract with increasing probability over time^62^. Our data are fully consistent with this concept, demonstrating the early generation and later maintenance of stem-like T cells with an early differentiation profile. Simultaneously, we observed an acute accumulation of “hot” effector subset cells with an IFN response signature, which were later replaced by “cold” effector subset cells that were devoid of this signature.

A central question for vaccinology, but also human T cell biology, is whether and how TCR avidity shapes the repertoire evolution of vaccination-induced T cells. Generally, we saw a higher degree of clonality at acute compared to memory time points, with diverse repertoires of TCRs being maintained nonetheless. Intuitively and based on studies using mouse models, we expected clonal expansion and maintenance to be driven by maximum TCR avidity. We were therefore surprised to observe dynamic clonal replacement patterns, and that the most prominently maintained and expanded clones did not show the highest functional avidities.

To understand these seemingly counterintuitive findings, it is important to clarify what is meant by “high” and “low” TCR avidity. We and others could previously show that the naïve repertoire encompasses TCRs with massive avidity differences^14,68,69^. Our here presented data on the functionality of vaccination-induced human antigen-specific T cell repertoires clearly support a significant enrichment of high-functionality TCRs in the antigen-experienced repertoire, while clonotypes having an EC_50_ value of less than -log_10_ 5 M are retained in a naïve phenotypic state. Our human data thereby, for the first time, confirm findings obtained by us and others in mouse models that showed efficient recruitment of high-avidity TCRs and variable or no recruitment of low-avidity TCRs^14,70,71^. Thus, the long-standing concept of TCR “avidity maturation” is valid on the population level and in the sense that immunization leads to clonal expansion and maintenance of high-functionality TCRs.

When there are only nuanced differences in avidity (e.g., EC_50_ varying between -log_10_ 6 M to 7 M for LTD), TCRs with maximum avidity did not dominate antigen-experienced repertoires that were of high overall functionality. Instead, we observed clonal expansion mostly for TCRs with “intermediate-high” avidities (EC_50_ of about -log_10_ 6.5 M), which could be explained by a higher number of clonotypes with such features compared to “low-high” or “high-high” avidity clonotypes. Why could the antigen-experienced repertoire be enriched for clonotypes with “intermediate-high”, but not “low-high” or “high-high” avidities? “High-high” avidity requires optimal structural solutions of TCR-pMHC recognition. It is intuitive that such optimal solutions are not frequently generated. In contrast, a lack of higher numbers of “low-high” clones may be explained by the fact that initial recruitment into antigen-experienced responses becomes less likely once a certain avidity threshold is not reached.

Many mouse studies that previously investigated the influence of TCR avidity on T cell fate used adoptive transfers of monoclonal cell populations with defined avidities. Usually, TCRs with large functionality differences are thereby chosen to reveal “deterministic” effects of TCR avidity. However, endogenous natural T cell responses such as those induced by vaccination actually develop from naïve precursor repertoires that are extremely diverse, with most TCRs being represented by a single cell^14,72^. Single-cell derived T cell responses are particularly susceptible to stochasticity^73–75^, possibly owing to a differential spatiotemporal reception of TCR signals 1, 2, and 3 during priming^66,76^. For clonotypes with the same TCR, the progeny size can vary 100.000-fold^75^. In line with such stochastic clonal expansion of individual clones, for many TCRs from our study we detected little or no clonal expansion at all even if their avidities were similar to those of the most expanded clones. Since clonal expansion was more likely, however, for clones with a more frequently observed avidity range (“intermediate-high”), we consider the term “probabilistic” most accurate.

Our study is restricted to the human T lymphocyte response in peripheral blood. Future work will need to investigate the influence of TCR avidity on human T cell fate also in secondary lymphoid and mucosal tissues. While we present a detailed investigation of the complete repertoires of five donors encompassing two HLA-I restricted epitope specificities with high immunogenicity, it remains to be elucidated whether clonal selection processes may follow different rules for less immunogenic epitopes.

What could be a physiological advantage of maintaining a diverse repertoire with overall high TCR functionality? Polyclonal repertoires are more flexible in counteracting mutational escape of epitopes^14^. Accordingly, we observed varying functionality of individual TCRs against mutations of the LTD epitope, with only polyclonal repertoires ensuring recognition of peptide alterations in a robust manner. One could speculate that a more deterministic effect on clonal expansion through maximum TCR avidity could lead to an undesired skewing of the TCR repertoire that optimizes recognition of a current epitope version, but would come at the cost of decreased recognition robustness of unpredictable mutant epitopes.

In terms of clinical translation, our findings therefore suggest that vaccination-induced T cell responses should harbor diverse clonotypes (ideally more than a dozen; **Fig. 5G**) with sufficiently high, but not necessarily “maximum” avidities. Given the importance of high TCR functionality of individual clones and polyclonality of the population as a whole, these two dimensions of antigen-specific T cell responses represent so-far neglected quality criteria for the assessment of vaccine immunogenicity. Fine-mapping of vaccination-induced T cell qualities is particularly relevant for the development of vaccines against pathogens like HIV, malaria, or tuberculosis, for which functional T cell responses are considered to be of special importance^1^.

## Methods

### Study cohort CoVa-Adapt

Ethics approval was granted by the local Ethics Committee of the Medical Faculty of the University Hospital of Erlangen, Friedrich-Alexander University Erlangen-Nürnberg, Germany (350_20B). Samples were collected after informed consent of the donors. Analyses from this cohort have been previously published^43^ and the cohort is described in detail in **Table 1** and **Suppl. Table 1**, HLA-typing was previously published^54^. In brief, donors were 23-58 years old (median: 42; interquartile range (IQR) 29-51), 55% female, of European Caucasian ethnicity, overall healthy (no chronic medication), of normal weight, and received a three-dose mRNA vaccination regimen with Comirnaty. Time intervals between immunizations were 22-28 days (median 23, IQR 23-25) between 1^st^ and 2^nd^ dose, and 196-244 days (median 227, IQR 216.5-229) between 2^nd^ and 3^rd^ dose. Blood was sampled before the 1^st^ dose, 8-13 days post 1^st^ dose (median 10, IQR 9-10), 9-11 days post 2^nd^ dose (median 10, IQR 10-11), 195-211days post 2^nd^ dose (median 210, IQR 209-210), 9-12 days post 3^rd^ dose (median 10, IQR 10-10) and 157-202 days post 3^rd^ dose (median 189, IQR 189-190). Additional blood samples were collected for eight donors 56-69 days (median 68, IQR 65.75-69) after 2^nd^ dose and for seven donors 101-126 days (median 108, IQR 108-113.5) after 3^rd^ dose. Some donors experienced SARS-CoV-2 breakthrough infections after day 10 post 3^rd^ dose which is indicated in the figures or figure legends if applicable. These breakthrough infections were self-reported and confirmed by PCR tests. 44% of breakthrough infections could be additionally validated by SARS-CoV-2 nucleocapsid^+^ serology, while no infection-free donor had detectable SARS-CoV-2 nucleocapsid-specific antibodies.

### Serum collection

Serum was isolated from venous blood collected in S-monovettes (Sarstedt, 227632). Samples were centrifuged for 10 min at 1200 rpm, 20°C, the serum supernatant was transferred into a new collection tube and centrifuged at maximum speed for 10 min in a tabletop centrifuge. Supernatants were collected again and stored at -20°C.

### Flow cytometry-based antibody assay

A previously published flow cytometry-based assay was used to detect spike-specific IgG, IgA and IgM antibodies^77^ as well as the different IgG subclasses^43^. In brief, stably transduced HEK293T cells with doxycycline-dependent expression of SARS-CoV-2 spike protein derived from the wild type strain were used as target cells. Spike protein-specific IgG, IgA, and IgM antibodies were quantified using a standard serum as described before^43^. For the binding assay, spike protein expression was induced by doxycycline treatment for 48 h, before 1×10^5^ cells were incubated with serum samples at various dilutions in 100 μL FACS-PBS (PBS with 0.5% BSA and 1 mM sodium azide) for 20 min at 4°. After washing, bound antibodies were detected via flow cytometry using anti-hIgG-AF647 (BioLegend, 410714), anti-hIgM-BV711 (BioLegend, 314540), anti-hIgA-FITC (SouthernBiotech, 2050-02).

Samples were measured on an AttuneNxt (ThermoFisher) and analyzed using FlowJo software 10.8.1 (Tree Star Inc.) and Graph Pad Prism9 (GraphPad Software). All sera with MFI values below the lowest limit of quantification (15.8 μg/mL for IgG; 15.6 FU/mL for IgA and IgM) were set to “n.d.”.

### Peripheral blood mononuclear cell (PBMC) isolation

PBMCs were isolated from citrated peripheral blood by density gradient centrifugation using a BioColl density medium with a density of 1.077 g/mL (BioSell, BS.L 6115). Cells were resuspended in heat-inactivated FCS + 10% DMSO and stored in liquid nitrogen.

### Multimerization of peptide-human leukocyte antigen (pHLA) monomers

Biotinylated HLA-A*01:01 molecules loaded with LTDEMIAQY-peptide (SARS-CoV-2 spike protein) or biotinylated HLA-A*02:01 molecules loaded with YLQPRTFLL-peptide (SARS-CoV-2 spike protein) were generated according to standard protocols of the laboratory of Dirk H. Busch, Munich, as describe before^78^. pHLA monomers were multimerized on a streptavidin backbone conjugated with PE-fluorophores (Life Technologies, 12-4317-87). Per 1×10^6^ cells, 0.2 μg of pHLA molecules were mixed with 0.125 μg of streptavidin-PE in 25 μL of FACS-buffer (PBS + 0.5% BSA) and incubated on ice for 30 min, unless specified otherwise. pHLA-multimers were directly used to stain cells for flow cytometry (see below).

### Multiparametric flow cytometry for T cells

The following antibodies were used for T-cell analysis: from BD Biosciences: anti-CD4-BV711 (740769, 1:400), anti-CD4-BUV395 (564724, 1:200), anti-CD19-PE/CF594 (562294, 1:200), anti-IFNγ-FITC (340449, 1:10), anti-IL-2-APC (341116, 1:20); from BioLegend: anti-CD8-APC (301049, 1:400), anti-CD4-BV510 (300545, 1:50), anti-human a/b TCR (hTCR)-PE (306708, 1:200), anti-human a/b TCR (hTCR)-FITC (306706, 1:200), anti-mouse TCR β chain (mTRBC)-APC/Fire^TM^750 (109246, 1:100); from eBioscience: anti-CD4-PE (12-0049-42, 1:400), anti-CD8-eF450 (48-0086-42, 1:200), anti-CD56-FITC (11-0566-42, 1:200), anti-CD45-PerCP/Cy5.5 (45-0459-42, 1:100), anti-CD45-PE/Cy7 (25-9459-42, 1:400); from Dako: CD45-PacificBlue (PB986, 1:50). For the viability staining, ethidium-monoazide-bromide (EMA) (ThermoFisher, E1374), propidium iodide (PI) (ThermoFisher, P1304MP), the ZombieAqua, or ZombieNIR Fixable Viability Kit (BioLegend, 423101 or 423102, 1:500) were used.

PBMCs or TCR-engineered T cells were first washed twice with cold FACS buffer. If required, pHLA-multimers were added (25 μL per sample with 1×10^6^ cells) and cells were incubated on ice for 25 min. Then, 25 μL of surface antibody mix including viability staining were directly added and cells were incubated on ice for additional 20 min. If no staining with pHLA-multimers was performed, cells were directly resuspended in 25 μl of surface antibody mix including viability staining and incubated on ice for 20 min. Staining with surface antibodies was followed by two washing steps with cold FACS buffer. Where applicable, cells were fix-permeabilized with the BD Cytofix/Cytoperm Kit (BD Bioscience, 554714) following manufacturer’s instructions. For intracellular cytokine staining, cells were incubated for 30 min on ice in 25 μL PermWash (1x) per sample with the respective antibodies. Cells were washed two times with cold PermWash (1x) and one time with cold FACS-buffer. Samples were acquired on a LSRFortessa Cell Analyzer (BD Biosciences) or a Northern Lights™ (Cytek Biosciences) and analyzed with FlowJo 10.7.2 (Tree Star Inc.).

### Peptide-induced cytokine expression by PBMCs

Cryopreserved PBMCs were thawed and rested overnight at 1×10^6^ cells/ml in complete RPMI medium (cRPMI: RPMI 1640 Medium + 10% heat-inactivated FCS, 0.05 mM β-mercaptoethanol, 0.05 mg/mL gentamicin, 1.1915 g/L HEPES, 0.2 g/L L-glutamine, 100 U/mL Penicillin-Streptomycin). 1×10^6^ PBMCs were stimulated with 11aa overlapping 15-mer PepMix^TM^ SARS-CoV-2 spike glycoprotein peptide pool (JPT, PM-WCPV-S-2) at a concentration of 1 μg/mL. Stimulation was performed for 20 h at 37°C in the presence of 1 μL/mL GolgiPlug™ (BD Biosciences, 555029). For the unstimulated condition, PBMCs were cultured in cRPMI medium and respective dilution of solvent DMSO. As a positive control, PBMCs were stimulated with 25 ng/mL phorbol myristate acetate (PMA) (Sigma-Aldrich, P1585-1mg) and 1 μg/mL ionomycin (Sigma-Aldrich, I9657-1MG). Following the stimulation, cells were stained with EMA for live/dead discrimination, followed by surface antibodies (CD4-PE, CD8-eF450) and intracellular cytokine staining (IFNγ-FITC, IL-2-APC). Samples were acquired on a LSRFortessa Cell Analyzer (BD Biosciences) and analyzed with FlowJo 10.7.2 (Tree Star Inc.).

### IFNγ Enzyme-linked immunospot (ELISpot)

Cryopreserved PBMCs were thawed and rested overnight at 1×10^6^ cells/ml in cRPMI medium. ELISpot plates (Millipore, MSIPS4510) were coated with anti-human IFNγ monoclonal antibody (clone 1-DIK, Mabtech, 3420-3-1000) at 0.5 μg/well overnight at 4°C. Plates were washed with sterile PBS and subsequently blocked with cRPMI medium for 1-2 h at 37°C. PBMCs were seeded at a density of 400,000 cells/well and stimulated with 11aa overlapping 15-mer PepMix^TM^ SARS-CoV-2 spike glycoprotein peptide pool or SARS-CoV-2 nucleoprotein peptide pool at a concentration of 1 μg/mL for 20 h at 37°C. For the unstimulated condition, PBMCs were cultured in cRPMI medium and respective dilution of solvent DMSO. As a positive control, PBMCs were stimulated with 25 ng/mL PMA and 1 μg/mL ionomycin. The following steps were performed at room temperature. Plates were washed with PBS containing 0.05% Tween-20 (Sigma-Aldrich, P9416-50ml) and incubated with biotinylated anti-human IFNγ monoclonal antibody (clone 7-B6-1, Mabtech, 3420-6-250) at 0.2 μg/well for 2 h. Plates were washed a second time with PBS containing 0.05% Tween-20 and subsequently incubated with an avidin-biotinylated peroxidase complex (VECTASTAIN^®^ Elite ABC-HRP Kit, Vector Laboratories, VEC-PK-6100) for 1-2 h. Afterwards, plates were washed first with PBS containing 0.05% Tween-20 following one washing step with PBS. Plates were developed by the addition of AEC substrate solution (Sigma-Aldrich, 152224-10ml) for 15 min, washed with water, and dried for 24 h in the dark. Analysis was performed on an ImmunoSpot^®^ Analyzer (Cellular Technologies Limited).

### Single-cell RNA sequencing of T cells

Single-cell RNA sequencing was performed for PBMCs from 13 ‘CoVa-Adapt’ donors at a total of seven time points after primary (P), secondary (S), and tertiary (T) vaccination as well as for PBMCs from HIM 189 days after 215^th^ vaccination (see **Suppl. Table 2**). PBMCs were thawed from cryopreserved stocks and rested overnight at 1×10^6^ cells/mL in cRPMI medium.

Experiment 1 and 2: Dextramer cocktails were prepared immediately before staining. Each donor was stained with all SARS-CoV-2 spike dextramers irrespective of HLA-match (see **Suppl. Table 2**). Per 5×10^6^ cells, 1 μl of HLA-matched dCODE dextramers^®^ (Immudex) and 0.2 μl 100 μM d-biotin (per dCODE dextramer^®^ specificity) were pre-mixed in FACS buffer to block free binding sites. 5×10^6^ PBMCs per donor and time point were recovered and first stained with a cocktail of surface antibodies (anti-CD19-PE/CF594, anti-CD56-FITC, anti-CD8-APC) including individual anti-CD45 antibodies (anti-CD45-PacificBlue, anti-CD45-PerCP/Cy5.5, anti-CD45-PE/Cy7) and individual hashtag antibodies (2.5 μL per 5×10^6^ PBMCs of TotalSeq-C anti-human hashtag antibodies 1–8, BioLegend, 394661, 394663, 394665, 394667, 394669, 394671, 394673, 394675) for 30 min on ice. Cells were washed twice with cold FACS-buffer and up to 8 samples were pooled. Pooled samples (40×10^6^ cells) were stained with prepared dextramer pools in a total volume of 400 μl for 30 min on ice. Cells were washed four times with cold FACS-buffer and live/dead discrimination was performed using PI immediately before sorting. Pooled samples could be distinguished by individual CD45 colour-barcoding at the sorter and by individual TotalSeq-C anti-human hashtag antibodies in the scRNAseq dataset. Single, live, CD19- and CD56-negative, CD8-positive, dextramer-positive lymphocytes were sorted in previously FCS-coated 1.5 mL tubes filled with FACS-buffer. Cells were sorted on a FACS Aria II cell sorter (BD).

Experiment 3: For each donor, individual dextramer cocktails were prepared directly before staining (see **Suppl. Table 2**). For each cocktail, 1 μL per 5×10^6^ cells of HLA-matched dCODE dextramers^®^ (immudex) and 100 μM d-biotin (1/10 of total dextramer volume) were pre-mixed in FACS-buffer to block free binding sites. 5×10^6^ PBMCs per donor and time point were recovered and first stained with 50 μL of individual dextramer cocktails for 30 min on ice. Afterwards, a cocktail of surface antibodies and viability staining (anti-CD19-PE/CF594, anti-CD56-FITC, anti-CD8-APC, anti-CD4-BV510, ZombieNIR), individual anti-CD45 antibodies (anti-CD45-PacificBlue, anti-CD45-PerCP/Cy5.5, anti-CD45-PE/Cy7), individual hashtag antibodies (2.5 μL per 5×10^6^ PBMCs of TotalSeq-C anti-human hashtag antibodies 1–8, BioLegend, 394661, 394663, 394665, 394667, 394669, 394671, 394673, 394675), and TotalSeq-C antibodies (0.078 μg per 5×10^6^ PBMCs anti-human CD45RA (BioLegend, 304163), 0.277 μg per 5×10^6^ PBMCs anti-human CCR7 (BioLegend, 353251), 0.25 μg 5×10^6^ PBMCs anti-human CXCR3 (BioLegend, 353251)) were added. For three samples, instead of individual TotalSeq-C antibodies, the TotalSeq-C Human Universal Cocktail, V1.0 (BioLegend, 399905) was added. On the day of the experiment, three vials of this universal cocktail were rehydrated in 18 μL of FACS buffer each, incubated for 5 min at room temperature, vortexed, and centrifuged at maximum speed for 30 sec at room temperature. One vial was used to stain 5×10^6^ PBMCs. Samples were incubated for an additional 30 min on ice. Cells were washed four times with cold FACS buffer and up to 8 samples were pooled prior to the sort. Pooled samples could be distinguished by individual CD45 colour-barcoding at the sorter and by individual TotalSeq-C anti-human hashtag antibodies in the scRNAseq dataset. Single, live, CD19-and CD56-negative, CD4-or CD8-positive, dextramer-positive lymphocytes were sorted in previously FCS-coated 1.5 mL tubes filled with FACS buffer. Additionally, single, live, CD19-and CD56-negative, total CD4- or CD8-positive lymphocytes were sorted from three donors, as a framework for CD4-and CD8-T-cell phenotypic clusters. Cells were sorted on a FACS Aria II cell sorter (BD).

Immediately after sorting, cells were loaded to a Chromium Next GEM Chip K (10X Genomics) and Chromium Next GEM Single-Cell 5′ kits v.2 were used to generate GEX, VDJ and CITEseq libraries according to the manufacturer’s instructions (10X Genomics, 1000263, 1000256, 1000252, 1000286, 1000250, 1000215, 1000190). Libraries were sent to Novogene (Cambridge, UK) and sequenced on an Illumina NovaSeq platform with PE150 strategy.

### Computational single-cell RNA sequencing data analysis

All sequencing runs were separately processed by the cellranger ‘multi’-command (version 6.0.2, 10x Genomics) with the gene expression reference GRCh38 (version 2020A, 10x Genomics), the VDJ reference vdj-GRCh38 (version 5.0.0, 10x Genomics), and customized feature barcode references. The remaining single-cell processing was conducted via Scanpy (v.1.8.2)^79^ and Scirpy (v.0.10.1)^80^ mainly as described by Heumos et al.^81^.

The resulting GEX and antibody capture count matrices were combined with the TCR contig annotation for each sequencing run individually. Each run was filtered for doublet and dying cells by run-specific thresholds on minimal (Experiment 1: 1000; Experiment 2: P10 sample: 1500, S10 sample: 1300, S210 sample: 1300, T10 sample: 1100; Experiment 3: sample1: 1200, sample2: 1000, sample3: 2500) and maximal (Experiment 1: 7000; Experiment 2: P10 sample: 9500, S10 sample: 8000, S210 sample: 8000, T10 sample: 9000; Experiment 3: sample1: 13000, sample2: 8500, sample3: 6000) UMI counts, minimal number of detected genes (Experiment 1: 1000; Experiment 2: P10 sample: 800, S10 sample: 700, S210 sample: 700, T10 sample: 650; Experiment 3: sample1: 750, sample2: 500, sample3: 1000), and a maximal fraction of mitochondrial genes (10% for all experiments). Additionally, genes expressed in less than ten cells were removed. The GEX data were normalized to 10,000 counts per cell and log1p-transformed. The sequencing runs were separated into donors and time points based on their hashtag antibody counts through HashSolo^82^ filtering additional doublet and unannotated cells. Afterwards, the samples were concatenated and all cells without annotated TCR were removed. Surface protein counts were transformed by the centered-log-ratio over the whole dataset.

Clonotypes were defined by having identical a- and β-CDR3 amino acid sequences on the primary or secondary chain. Clonal expansion was calculated across the dataset as well as for each donor and time point, individually. Cell-level scores were calculated through gene sets described by Seumois^52X^ and Szabo^51^. Dextramer specificity was assigned if the dextramer UMI counts surpassed an experiment- and epitope-specific threshold (see **Suppl. Table 3**) and represented at least 40% of the total UMI counts of this cell. Additionally, specificity was only assumed if 50% of a clonotype’s cells were assigned specific (after UMI and per-cell purity cut-off) and the donor matched the dextramers HLA. For the assignment, the dextramers KVFRSSVLH, RLFRKSNLK, and GTHWFVTQR were excluded due to a high level of unspecific staining. CD8 and CD4 T cells were identified based on the gene expression level on *CD4, CD8A*, and *CD8B* (threshold: 0.5) and the surface proteins Hu.CD8 (threshold: 0.75) and Hu.CD4_RPA.T4 (threshold: 0.95). CD8 T cells were defined by elevated *CD8A*-, *CD8B*-gene expression, or CD8 surface protein markers, while cells without these or additional CD4 gene or protein markers were removed. Ambiguous or unannotated cells were also removed from the dataset.

Analysis was performed only on CD8 T cells using the 5,000 highly variable genes excluding TCR-forming genes. UMAP visualization^83^ was calculated at 15 neighbors and Leiden clustering^84^ at a resolution of 0.75. Differentially expressed genes and surface proteins were calculated via a t-test with Benjamini-Hochberg correction using the ‘scanpy.tl.rank_genes_groups’ function. Diffusion pseudotime^85^ was estimated through ‘scanpy.tl.dpt’ using the cell with the smallest second diffusion map component as root. Diffusion pseudotime was calculated over all cells (**Fig. 2F**) but mainly visualized for conventional T cell clusters. Distances between clonotype sequences were calculated via TCRdist3^86^. Following Pogorelyy et al.^87^, connectivity between clonotypes was defined at a threshold not exceeding 120 and clones within the top 1% centrality were removed. The resulting graph was visualized using the ‘Circle Pack Layout’ with Connected components and Modularity at standard settings as Hierarchy in Gephi (v.0.9.7)^88^.

### Data and Code Availability

All single-cell sequencing data are available and publicly accessible at NCBI GEO under the accession numbers GSE261966, GSE261967, and GSE249998. A processed and annotated version of the data can be accessed at Zenodo (10.5281/zenodo.13981508). The source code to analyze the sequencing data is available on GitHub at https://github.com/SchubertLab/CovidVac_CD8.

### Transgenic TCR re-expression in primary human T cells

A coherent description for the workflow of targeted TCR re-expression in primary human T cells using CRISPR-Cas9-mediated orthotopic TCR replacement has previously been published^57,59^. A brief description including all relevant alterations to the published protocol are summarized in the following chapters ‘Transgenic TCR DNA template design’, ‘Double-stranded DNA production’, ‘T cell activation for genetic editing’, ‘Ribonucleoprotein production’, and ‘Orthotopic TCR replacement (OTR)’.

### Transgenic TCR DNA template design

The DNA template was designed *in silico* and synthesized by Twist Bioscience. The construct had the following structure: The left homology arm (LHA; 396 bp) was followed by a self-cleaving peptide P2A and the TCR β-chain which consisted of the human variable part (VDJβ) and the murine TCR β constant region with an additional cysteine bridge (mTRBC-Cys)^89^. The subsequent self-cleaving peptide T2A separated the β-chain from the following a-chain which was designed according to the same principle with the human variable part (VJa) being followed by the murine TCR α constant region with an additional cysteine bridge (mTRAC-Cys)^89^. After the stop codon (TGA) and the bovine growth hormone polyA signal (bGHpA), the 330 bp right homology arm (RHA) concluded the HDR template. For sequences of these segments see **Suppl. Table 5** and **Suppl. Table 6**.

### Double-stranded DNA production

The DNA construct was ordered as a sequence-verified plasmid gene via a commercial provider (Twist Bioscience). The lyophilized plasmid was reconstituted with sterile water to 60 ng/μL and amplified by PCR to generate a linearized double-stranded HDR template. A Cas9 Target Sequences (CTS) was incorporated at the 5’-end of the HDR template by the genomic primer targeting the *hTRAC* LHA. Each 100 μl PCR reaction contained 1 x Herculase II Reaction Buffer, 0.4 μM *hTRAC* HDR genomic forward primer targeting LHA (5’-TCTCTCTCTCAGCTGGTACACGGCTGCCTTTACTCTGCC AGAG-3’), 0.4 μM *hTRAC* HDR genomic reverse primer targeting RHA (5’-CATCATTGACCAGAGCTCTG-3’), 0.5 mM dNTPs, 1 μL Herculase II Fusion DNA Polymerase, and 60 ng reconstituted DNA in PCR grade water. The PCR was run with the following cycling conditions: Initial denaturation at 95°C for 3 min, 34 cycles of 95°C for 30 sec, 63°C for 30 sec and 72°C for 3 min, final elongation at 72°C for 3 min, and hold at 4°C. Successful amplification was confirmed with an 1% agarose gel and amplified HDR template was purified with a MinElute PCR Purification Kit (Qiagen, 28004) according to the manufacturer’s instructions.

### T cell activation for genetic editing

PBMCs were isolated from blood provided by healthy volunteers (Transfusion Medicine, University Hospital Erlangen) and frozen at -80°C for storage. Ethics approval was granted by the local Ethics Committee of the Medical Faculty of the University Hospital of Erlangen, Friedrich-Alexander University Erlangen-Nürnberg, Germany (392_20Bc). Samples were collected after informed consent of the donors. For T-cell activation, these PBMCs were thawed and overnight-rested at 2×10^6^ cells/mL in cRPMI medium supplemented with 50 U/mL Interleukin-2 (IL-2). Afterwards, PBMCs were activated for two days at 1×10^6^ cells/mL on tissue-culture plates with 1 μg/μl of surface-bound anti-CD3 (BioLegend, 317302) and anti-CD28-antibodies (BioLegend, 302902) in medium supplemented with 300 U/mL IL-2 (Peprotech, 200-02), 5 ng/mL Interleukin-7 (Peprotech, 200-07), and 5 ng/mL Interleukin-15 (Peprotech, 200-15).

### Ribonucleoprotein production

Activated PBMCs were electroporated with ribonucleoproteins (RNPs) targeting the endogenous *hTRAC* and *hTRBC* locus as well as the purified HDR template. For final electroporation, 3.5 μL of *hTRAC* and 3 μl of *hTRBC* RNP (final concentration 20 μM) were required per electroporation sample. First, 40 μM gRNAs were produced by mixing equimolar amounts of trans-activating crRNA (tracrRNA) (Integrated DNA Technologies, 1072534) with *hTRAC* crRNA (5’-AGAGTCTCTCAGCTGGTACA-3’^90^, Integrated DNA Technologies) or *hTRBC* crRNA (5’-GGAGAATGACGAGTGGACCC-3’^57^, Integrated DNA Technologies) and incubating the mixtures at 95°C for 5 min with subsequent cool down to room temperature. Afterwards, 50 μg/sample poly-L-glutamic acid (PGA; Sigma-Aldrich, P4761) were added to *hTRAC* gRNA^91,92^ and 20 μM electroporation enhancer (Integrated DNA Technologies, 10007805) were added to both *hTRAC* and *hTRBC* gRNA. RNP production was concluded by adding equal volume of Cas9 Nuclease V3 (Integrated DNA Technologies, 1081059, diluted to 6 μM) to *hTRAC* and *hTRBC* gRNA (40 μM) respectively. RNPs were incubated for 15 min at RT and subsequently stored on ice for processing at the same day. For the calculations above, the volume of PGA was not considered.

### Orthotopic TCR replacement (OTR)

Prior to electroporation, DNA-sensing inhibitors RU.521 (small-molecule inhibitor of cyclic GMP-AMP synthase (cGAS); InvivoGen, inh-ru521)^91^ was added to the cells at a final concentration of 4.82 nM for 6 h. Afterwards, activation was stopped by transferring cells to a new plate in fresh cRPMI medium. For electroporation, 1×10^6^ activated cells per electroporation sample were resuspended in 20 μL P3 electroporation buffer (Lonza, V4SP-3960) and then mixed with DNA/RNP mix (0.5 μg HDR template, 3.5 μL *hTRAC* and 3 μL *hTRBC* RNPs). After transfer into the 16-well Nucleocuvette^TM^ Strip (Lonza, V4SP-3960), cells were electroporated (pulse sequence EH100) in the Lonza 4D-Nucleofector^TM^. After electroporation, cells were rescued in 180 μl of antibiotic-free RPMI medium supplemented with 180 U/mL IL-2. After 15 min, 20 μl of a mixture containing 0.5 μM HDAC class I/II Inhibitor Trichostatin A (AbMole, M1753) and 10 μM DNA-dependent protein kinase (DNA-PK) inhibitor M3814 (chemietek, CT-M3814) was added to each sample^93^. Cells were incubated for 12-18 h in a 96-well U-bottom plate, before the medium was supplemented with an antibiotic mix containing gentamicin, penicillin and streptomycin to produce cRPMI medium. 24 h after electroporation, cells were transferred into a 24-well plate and cultivated in a final volume of 1 ml fresh cRPMI medium supplemented with 180 U/ml IL-2. Four days after electroporation, successful editing was validated by flow cytometry (anti-CD4-PE, anti-CD8-eF450, anti-human a/b TCR (hTCR)-FITC, anti-mouse TCR β chain (mTRBC)-APC/Fire^TM^750, ZombieAqua). Cells were cultivated for additional seven to eight days in cRPMI with addition of 50 U/ml IL-2 every two days before specificity was determined by multimer staining and peptide-induced cytokine expression.

### K562 peptide pulsing

To assess reactivity of TCR-engineered T cells against the SARS-CoV-2 spike epitopes LTDEMIAQY (including altered peptide ligands (APLs) ATDEMIAQY, LADEMIAQY, LTAEMIAQY, LTDAMIAQY, LTDEAIAQY, LTDEMAAQY, LTDEMIAAY, LTDEMIAQA), YLQPRTFLL, and QYIKWPWYI, engineered T cells were co-cultured with peptide-pulsed K562 HLA-A*01:01^+^, HLA-A*02:01^+^or HLA-A*24:02^+^ target cells, respectively. The day before the co-culture, K562 cells were irradiated at 80 Gray and subsequently washed twice with cRPMI. Irradiated K562 cells were resuspended to a cell density of 3×10^6^ cells/mL and pulsed with LTDEMIAQY, YLQPRTFLL, or QYIKWPWYI-peptide ranging from 100 μM to 0.01 μM, if functional avidity of TCR-engineered T cells was determined. To assess cross-reactivity against potential escape mutations, irradiated K562 cells were resuspended to a cell density of 3×10^6^ cells/mL and pulsed with 10 μM of wild type LTDEMIAQY peptide or individual APLs. As a negative control, irradiated K562 cells were pulsed with respective dilution of solvent DMSO. Peptide-pulsed K562 cells were cultured overnight at 37°C.

### Peptide-induced cytokine expression by TCR-engineered T cells

The percentage of transgenic TCR (mTRBC)^+^ cells within the population TCR-engineered T cells was again determined the day before the co-culture. TCR-engineered T cells containing 5×10^4^ transgenic TCR (mTRBC)^+^ cells were co-cultured with 5×10^4^ peptide-pulsed K562 cells for 4 h at 37°C in cRPMI and in the presence of 1 μL/mL GolgiPlug™ (BD Biosciences, 555029). As a positive control, engineered T cells were stimulated with 25 ng/mL phorbol myristate acetate (PMA) (Sigma-Aldrich, P1585-1mg) and 1 μg/mL ionomycin (Sigma-Aldrich, I9657-1MG). Following the co-culture, cells were stained with surface antibodies including viability staining (panel LSRFortessa Cell Analyzer: anti-CD4-BUV395, anti-CD8-eF450, anti-CD19-PE/CF594, anti-human a/b TCR (hTCR)-PE, anti-mouse TCR β chain (mTRBC)-APC/Fire^TM^750, ZombieAqua; panel Northern Lights™: anti-CD4-BV711, anti-CD8-eF450, anti-CD19-PE/CF594, anti-human a/b TCR (hTCR)-PE, anti-mouse TCR β chain (mTRBC)-APC/Fire^TM^750, ZombieAqua) followed by intracellular cytokine staining (IFNγ-FITC, IL-2-APC). Samples were acquired on a LSRFortessa Cell Analyzer (BD Biosciences) or a Northern Lights™ (Cytek Biosciences) and analyzed with FlowJo 10.7.2 (Tree Star Inc.). To determine functional avidity, each TCR was tested in at least two independent experiments with two technical replicates each. Technical replicates were monitored between experiments and flow cytometers and excluded in case of large deviations. Dose-response curves and half-maximal cytokine production (IFNγ EC_50_) were calculated using Graph Pad Prism9 (GraphPad Software). Avidity on a population level was calculated weighing in clonal expansion as follows (with n=number of clones):

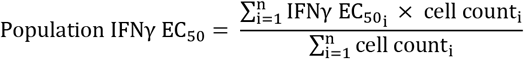

### Comparison of cell and clone count distributions

For A*01/LTD and A*02/YLQ specific TCR clones and specific cells detected at defined time points after vaccination, distributions were plotted as kde-plots. Similarity of the clone and cell count distribution was determined using a Kolmogorow-Smirnoff test. Analysis was performed using Python 3 including seaborn version 0.13.2 for plotting and script version 1.13.0 for statistics.

### Modelling of cumulative protection

To determine the protection against altered peptide ligands (APL) by certain TCR repertoires sizes, every TCR was first classified as protective or non-protective against every APL. The threshold for protection was defined by the reactivity of clone 297 against the wild type LTD peptide. Clone 297 showed the lowest avidity (Fig. 4C) that still led to active recruitment after vaccination as was therefore chosen for this threshold. Then every possible combination of TCRs for specific TCR repertoire sizes was tested for the degree of protection meaning for how many altered peptides protective reactivity is provided. Over all these permutations a curve representing the mean protection score with standard deviation was fitted. Analysis was performed using Python 3 including seaborn version 0.13.2 for plotting, itertools for generating the permutations and scipy version 1.13.0 for statistics.

## Author contributions

Conceptualization: K.S., data curation: K.K., F.D., formal analysis: K.K., F.D., M.G., F.G., funding acquisition: K.S., B.Sc. investigation: K.K., F.D., A.M.T., C.S., F.G., B.Sp., methodology: K.K., F.D., M.T., K.S., project administration: K.S., resources: E.D., D.H.B., C.B., M.T., B.Sc., K.S., software: F.D., B.Sc., supervision: B.Sc., K.S., validation: K.K., F.D., visualization: K.K., F.D., C.M., K.S., writing original draft: K.K., K.S., review & editing: all authors.

## Acknowledgments

This work is supported by grants from the German Federal Ministry of Education and Research (BMBF, projects 01KI2013 and 031L0290B), the Else Kröner-Fresenius-Stiftung (project 2020_EKEA.127) and the German Research Foundation (DFG) through the research training group RTG 2504 (project 401821119) to K.S.; the Interdisciplinary Center for Clinical Research (IZKF) at the University Hospital of the University of Erlangen-Nürnberg (advanced project A101) to M.T.; and the BMBF (project 031L0290A) to B.Sc.. F.D. is supported by the Helmholtz Association under the joint research school “Munich School for Data Science - MUDS” and acknowledges financial support from the Joachim Herz Stiftung. Funding agencies had no influence on the study design or implementation.

We thank members from the Schober laboratory and Veit R. Buchholz from Munich for experimental support and critical discussion. Furthermore, we also gratefully acknowledge the generous support by the Manfred Roth-Stiftung, Fürth, Germany, and thank the Core Unit Cell Sorting and Immunomonitoring Erlangen.

