## Supplementary Figures for "Quality of vaccination-induced T cell responses is conveyed by polyclonality and high, but not maximum, antigen receptor avidity"

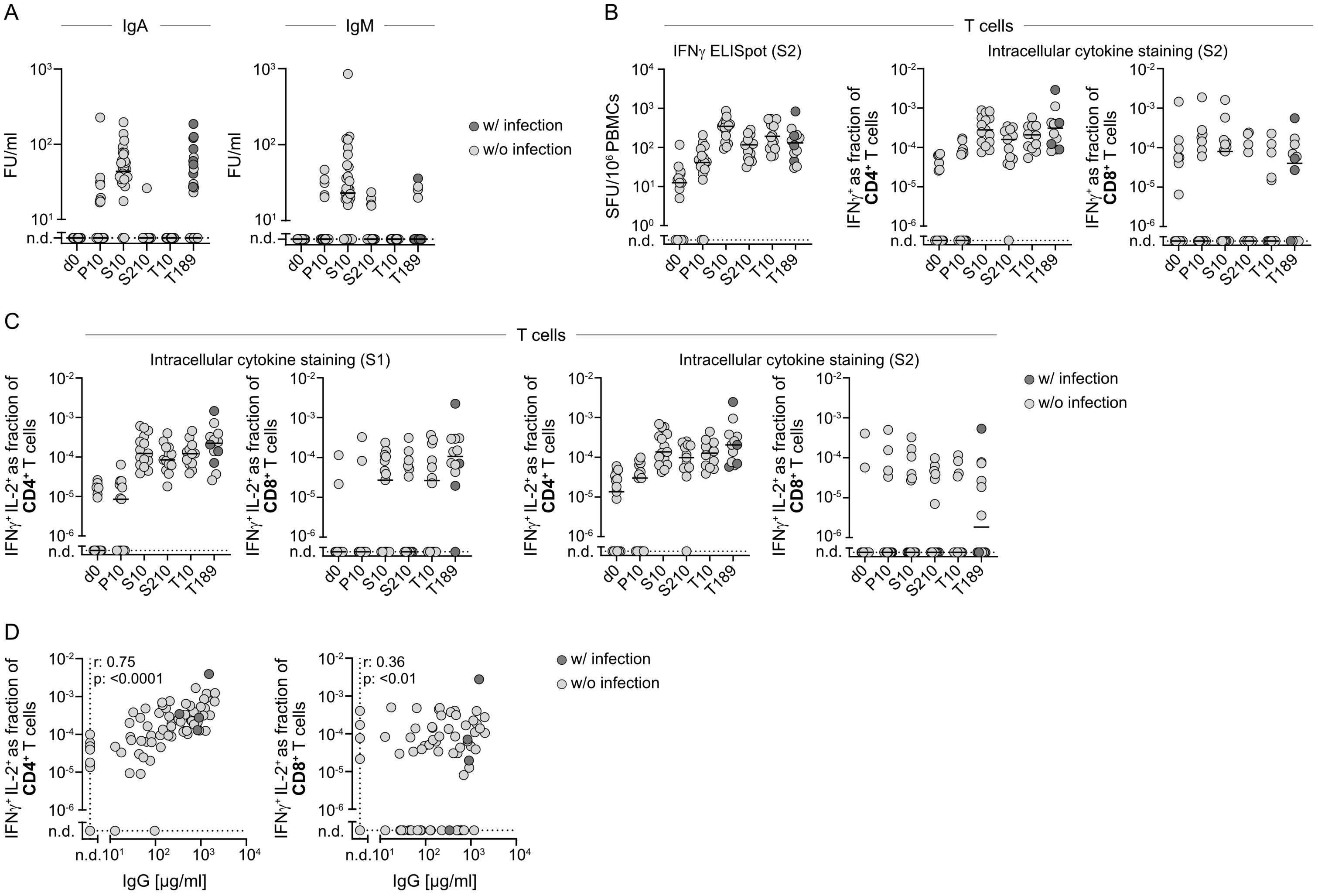


**Suppl. Fig. 1: Quantity of vaccination-induced antibody and T cell responses.** For all donors of the CoVa-Adapt study, serum and PBMCs were collected at day 0 (d0), 10 days after primary (P10), 10 and 210 days after secondary (S10, S210), and 10 and 189 days after tertiary (T10, T189) vaccination. Eleven donors experienced a breakthrough infection between T10 and T189. Donors with corresponding positive nucleocapsid serology are marked in dark grey in sub-figures A-D. **A** Spike-specific IgA and IgM in serum quantified by a flow cytometric assay using full-length spike protein as targets. Samples below the lower limit of quantification (15.6 FU/mL for IgA and IgM) were set to not detected (n.d.). Data points represent individual donors (n=27-29 per time point), solid lines indicate the mean. **B,C** Spike-specific CD4$^+$ and CD8$^+$ after 20h *in vitro* re-stimulation with 15mer peptides covering the complete wild type spike protein. **B** Peptides were provided in two sub-pools S1 (Fig. 1C,D) and S2 (Suppl. Fig. 1B). Quantification of spot forming units (SFU) for IFNγ ELISpot (left) and flow cytometric assessment of IFNγ$^+$ T cells as a fraction of living CD4$^+$ (middle) or CD8$^+$ (right) T cells. Data points represent individual donors (n=12-29 per time point), solid lines indicate the mean. Samples without IFNγ$^+$ T cells above the negative control were set to not detected (n.d.). **C** Quantification of IFNγ$^+$ IL-2$^+$ T cells measured by flow cytometry as a fraction of living CD4$^+$ or CD8$^+$ T cells after stimulation with peptide sub-pools S1 (left) and S2 (right). Data points represent individual donors (n=12-19 per time point), solid lines indicate the mean. Samples without IFNγ$^+$ IL-2$^+$ T cells above the negative control were set to not detected (n.d.). **D** Correlation of spike-specific IgG in serum with IFNγ$^+$ IL-2$^+$ T cells in flow cytometry (S1 and S2 stimulation combined) as a fraction of living CD4$^+$ (left) or CD8$^+$ (right) T cells. Samples with spike-specific serum IgG below the lower limit of quantification (15.8 µg/ml) or without IFNγ$^+$ IL-2$^+$ T cells above the negative control were set to not detected (n.d.). Data points represent individual donors across all time points (n=81). Correlation was determined by non-parametric spearman correlation.

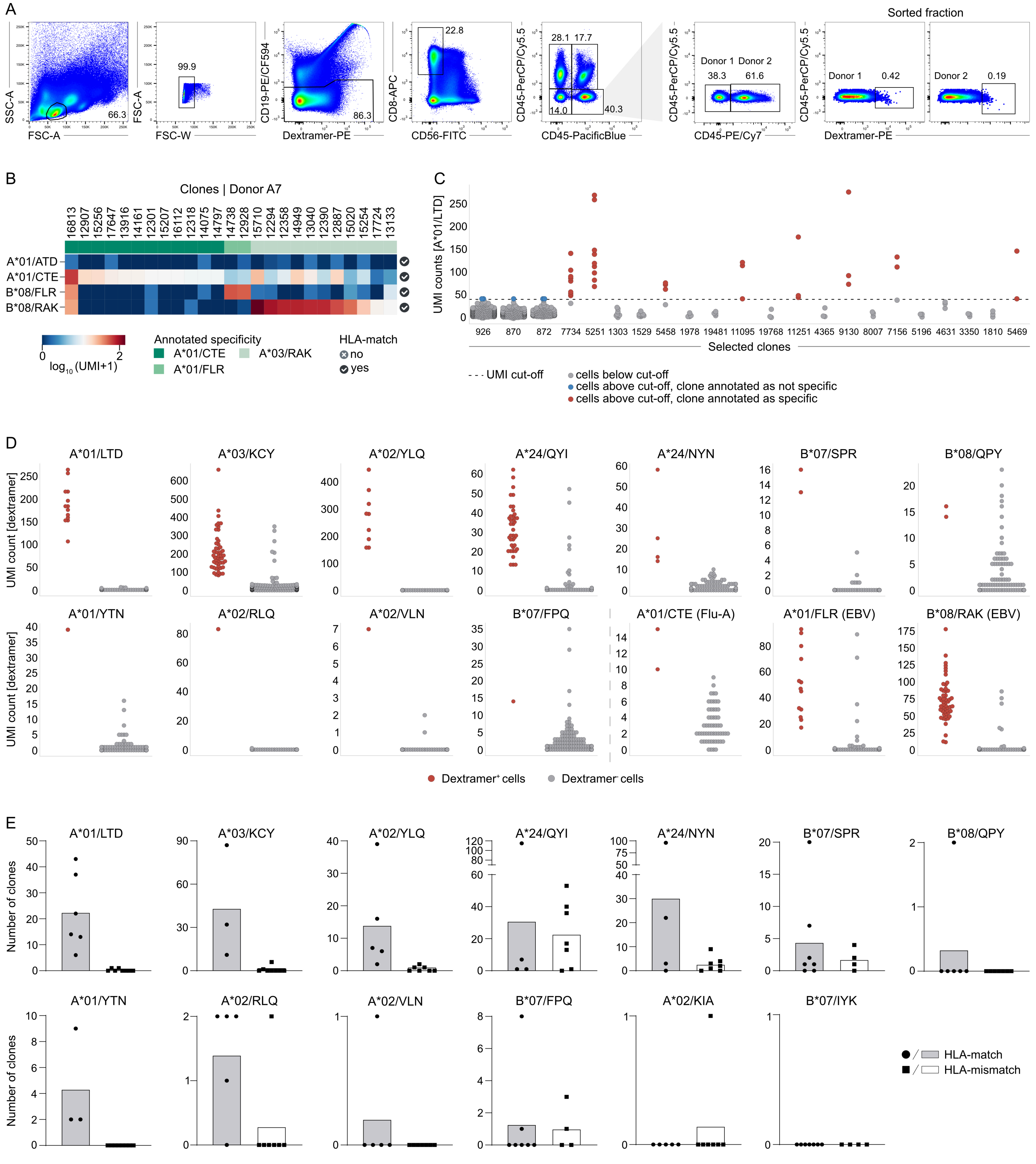


**Suppl. Fig. 2: Identification of SARS-CoV-2 spike epitope-specific CD8 T cells via dextramers. A** Representative gating strategy to enrich dextramer$^+$ T cells for scRNAseq via flow cytometric cell sorting. Single, live, CD19- and CD56-negative, CD8-positive, dextramer-positive lymphocytes were enriched for a total of 14 donors and seven time points after primary, secondary, and tertiary SARS-CoV-2 vaccination. Based on individual CD45 color-barcodes, cells from different donors were identified in the pooled sample to ensure balanced cell numbers per donor during sorting. During later analysis, donors were identified via hashtag antibodies. **B** Representative heatmap showing average UMI counts of detected clones with assigned epitope-specificity for dextramers of HHV-1 (A*01/ATD), Flu-A (A*01/CTE), and EBV (B*08/FLR, B*08/RAK) in sequencing experiment 3. CoVa-Adapt donor A7 is shown. For each epitope, donor-dependent HLA-matching is indicated. **C** Representative distributions of dextramer$^+$ and dextramer$^-$ cells from donor A8 of sequencing experiment 3. UMI counts for A*01/LTD dextramer are depicted for each cell of representative clones. UMI cut-off was set to 40 and depicted as a dotted line. Individual cells with UMI counts below this cut-off are depicted in grey. Cells with higher UMI counts are shown in blue if the overall clone was annotated as dextramer$^-$, and in red if the clone was assigned dextramer$^+$, after selection based on additional cell purity (40%) and clone purity (50%) criteria. **D** Distribution of dextramer UMI counts of cells annotated as dextramer$^+$ and dextramer$^-$. For each SARS-CoV-2 spike, Flu-A (A*01/CTE), and EBV (B*08/FLR, B*08/RAK) specificity, cells from one representative donor and one representative sequencing experiment are shown. **E** Number of SARS-CoV-2 spike dextramer$^+$ clones identified in HLA-matched (dots, grey bar) or HLA-mismatched (squares, white bar) donors over all three sequencing experiments. Data points represent individual donors, bars indicate the mean.

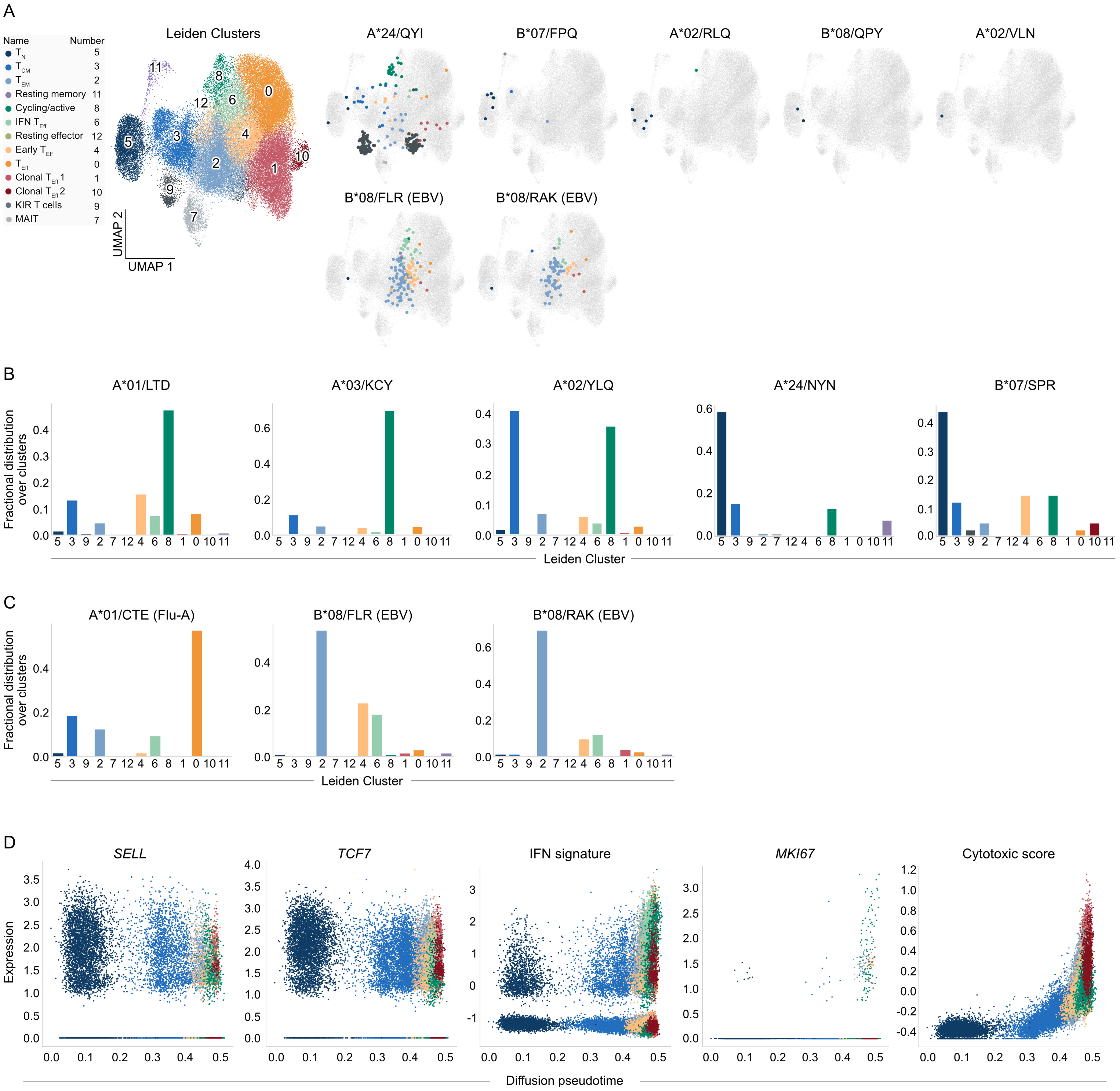


**Suppl. Fig. 3: Pooled phenotypes of epitope-specific CD8 T cells**. **A** UMAP with Leiden clusters (cluster numbers in UMAP, names and numbers on the left) of CD8$^{+}$ T cells enriched for dextramer-binding (Suppl. Fig. 2A) from three independent scRNAseq experiments (n = 53,907 cells). SARS-CoV-2 spike and EBV (B*08/FLR, B*08/RAK) epitope-specific T cells of all HLA-matched donors (A*24/QYI: n=4, B*07/FPQ: n=7, A*02/RLQ: n=5, B*08/QPY: n=6, A*02/VLN: n=5, B*08/FLR: n=3, B*08/RAK: n=3) and screened time points are visualized and colored based on their cluster annotation. Cells without the indicated epitope-specificity are shown in grey. **B, C** Fractional distribution of selected SARS-CoV-2 spike (B) and Flu-A (A*01/CTE) or EBV (B*08/FLR, B*08/RAK) (C) epitope-specific T cells over all Leiden clusters. Leiden Cluster are ordered based on diffusion pseudotime. **D** Log-normalized expression of selected genes and scores of individual cells. Cells are ordered by diffusion pseudotime and colored based on their cluster annotation. For the sake of clarity, expression is only shown for cells within conventional T cell clusters 5, 3, 2, 4, 6, 8, 1, 0, and 10.

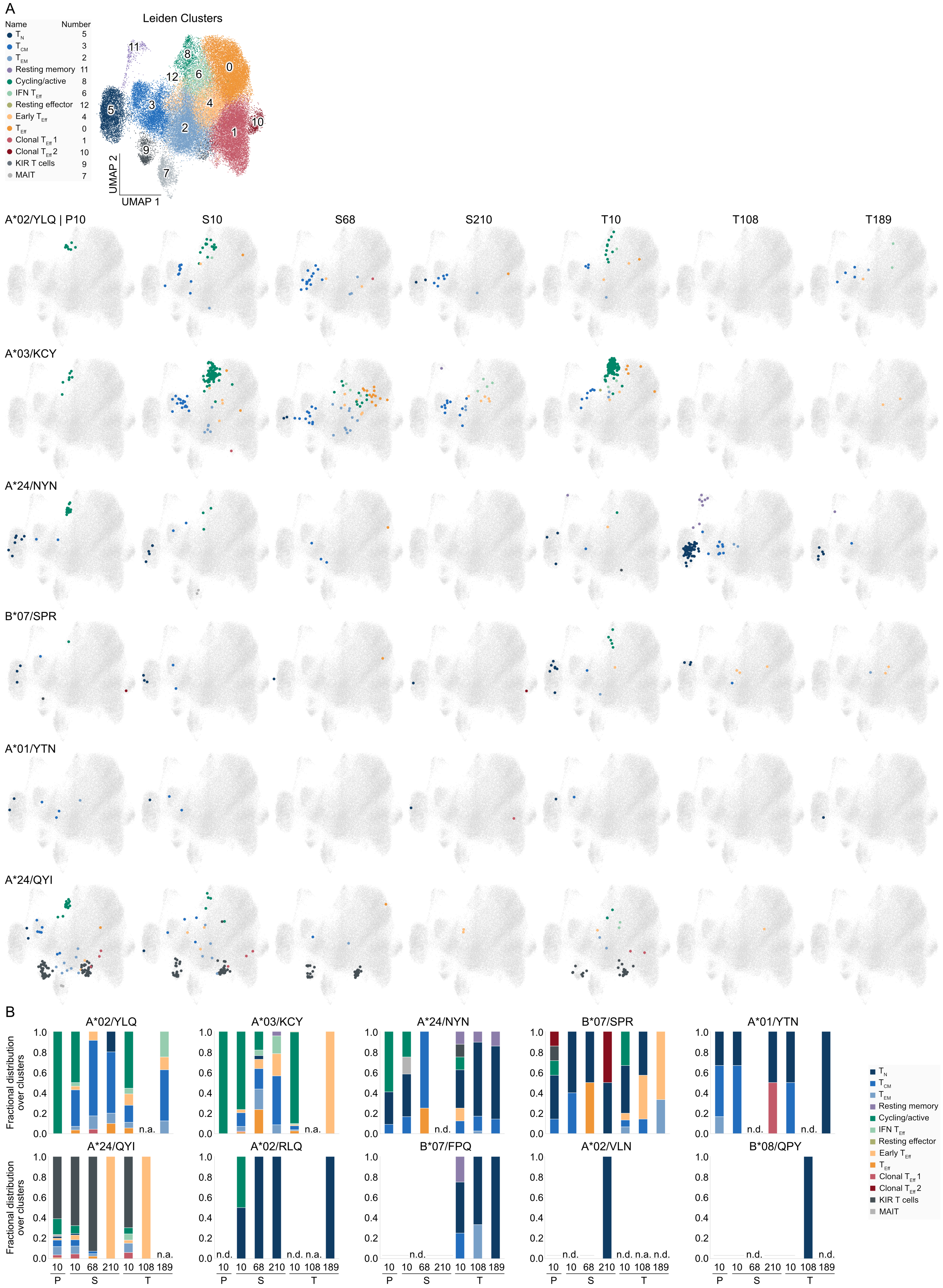

A
Leiden Clusters
Name
Number
T N 5
T CM 3
T EM 2
Resting memory 11
Cycling/active 8
IFN T Eff 6
Resting effector 12
Early T Eff 4
T Eff 0
Clonal T Eff 1 1
Clonal T Eff 2 10
KIR T cells 9
MAIT 7
UMAP 1
UMAP 2
A*02/YLQ | P10
S10
S68
S210
T10
T108
T189
A*03/KCY
A*24/NYN
B*07/SPR
A*01/YTN
A*24/QYI
B
A*02/YLQ
A*03/KCY
A*24/NYN
B*07/SPR
A*01/YTN
A*24/QYI
A*02/RLQ
B*07/FPQ
A*02/VLN
B*08/QPY
Fractional distribution over clusters
n.a.
n.d.
10 10 68 210 10 108 189
P S T

**Suppl. Fig. 4: Longitudinal phenotypes of SARS-CoV-2 spike epitope-specific CD8 T cells. A** UMAP with Leiden clusters (cluster numbers in UMAP, names and numbers on the left) of $CD8^+$ T cells enriched for dextramer-binding (Suppl. Fig. 2A) from three independent scRNAseq experiments (n = 53,907 cells). UMAP visualization of epitope-specific T cells of HLA-matched donors (A*02/YLQ: n=5, A*03/KCY: n=3, A*24/NYN: n=4, B*07/SPR: n=7, A*01/YTN: n=3, A*24/QYI: n=4) at individual time points in days after primary (P), secondary (S) and tertiary (T) vaccination. Epitope-specificities with detected cells at minimum five individual time points are shown. Colors represent cluster location of epitope-specific cells at respective time points. Cells without the indicated epitope-specificity are shown in grey. **B** Quantified fractional distribution over all clusters of epitope-specific T cells of HLA-matched donors (A*02/YLQ: n=5, A*03/KCY: n=3, A*24/NYN: n=4, B*07/SPR: n=7, A*01/YTN: n=3, A*24/QYI: n=4, A*02/RLQ: n=5, B*07/FPQ: n=7, A*02/VLN: n=5, B*08/QPY: n=6) at individual time points after vaccination. Colors represent cluster location of epitope-specific cells at respective time points. For each epitope-specificity, it is indicated whether no cell was detected at a certain time point after vaccination (n.d.) or no sample was acquired (n.a.).

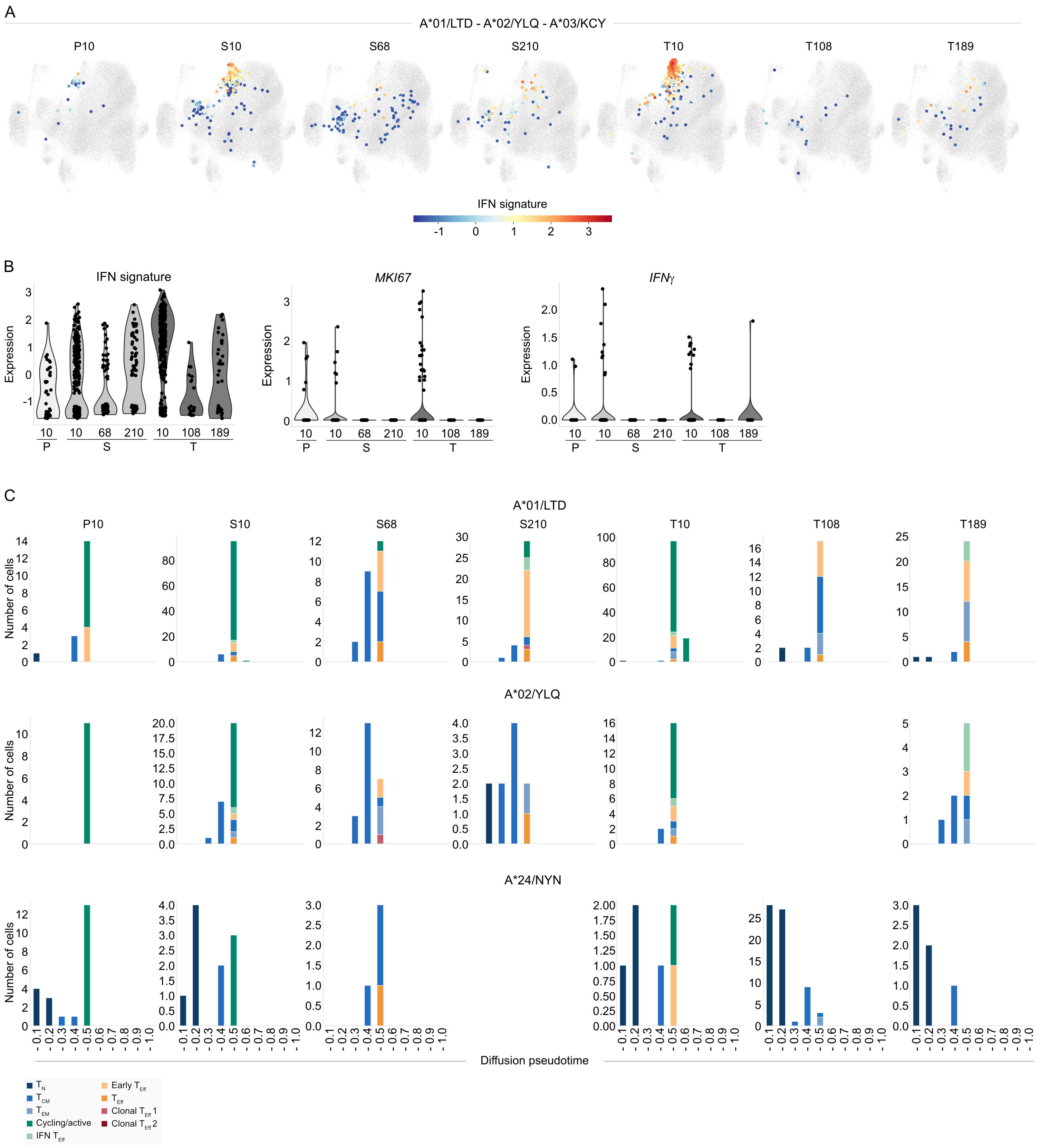


**Suppl. Fig. 5: Time point and differentiation-specific phenotypes of SARS-CoV-2 spike epitope-specific CD8 T cells. A** UMAP visualization of IFN signature expression at individual time points in days after primary (P), secondary (S) and tertiary (T) vaccination of A*01/LTD-, A*02/YLQ-, and A*03/KCY-specific T cells of HLA-matched CoVa-Adapt donors (n=13). **B** Log-normalized expression of selected genes and scores of individual A*01/LTD-, A*02/YLQ-, and A*03/KCY-specific T cells at individual time points after vaccination. **C** Number of A*01/LTD-, A*02/YLQ-, or A*24/NYN-specific T cells (y-axis) detected at individual time points after vaccination within a defined diffusion pseudotime interval (x-axis; -0.1 means 0.0-0.1, -0.2 means 0.1-0.2 etc.). Colors represent cluster location of epitope-specific cells. For the sake of clarity, only cells within conventional T cell clusters 5, 3, 2, 4, 6, 8, 1, 0, and 10 are shown.

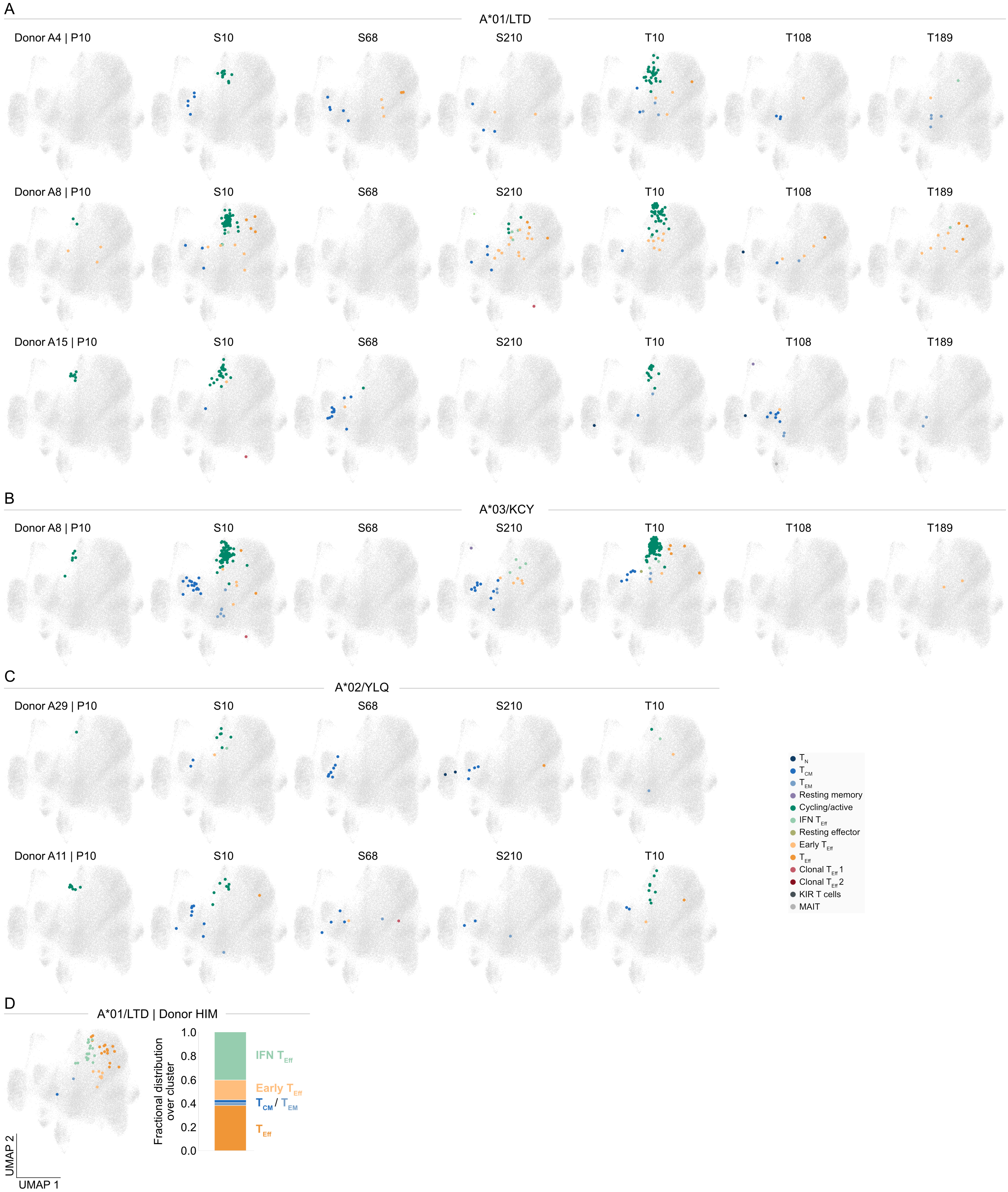


**Suppl. Fig. 6: Donor-specific longitudinal phenotypes of SARS-CoV-2 spike epitope-specific CD8 T cells. A-C** UMAP visualization of epitope-specific T cells of individual HLA-matched CoVa-Adapt donors at individual time points in days after primary (P), secondary (S) and tertiary (T) vaccination. A*01/LTD- (A), A*03/KCY- (B), and A*02/YLQ- (C) specific T cells are shown. Colors represent cluster location of epitope-specific cells at respective time points. Cells without the indicated epitope-specificity are shown in grey. **D** UMAP visualization (left) and quantified fractional distribution over all clusters (right) of A*01/LTD-specific T cells from donor HIM 189 days after 215 reported vaccinations (X189). Colors represent cluster location of A*01/LTD-specific cells. Cells without the indicated epitope-specificity are shown in grey.

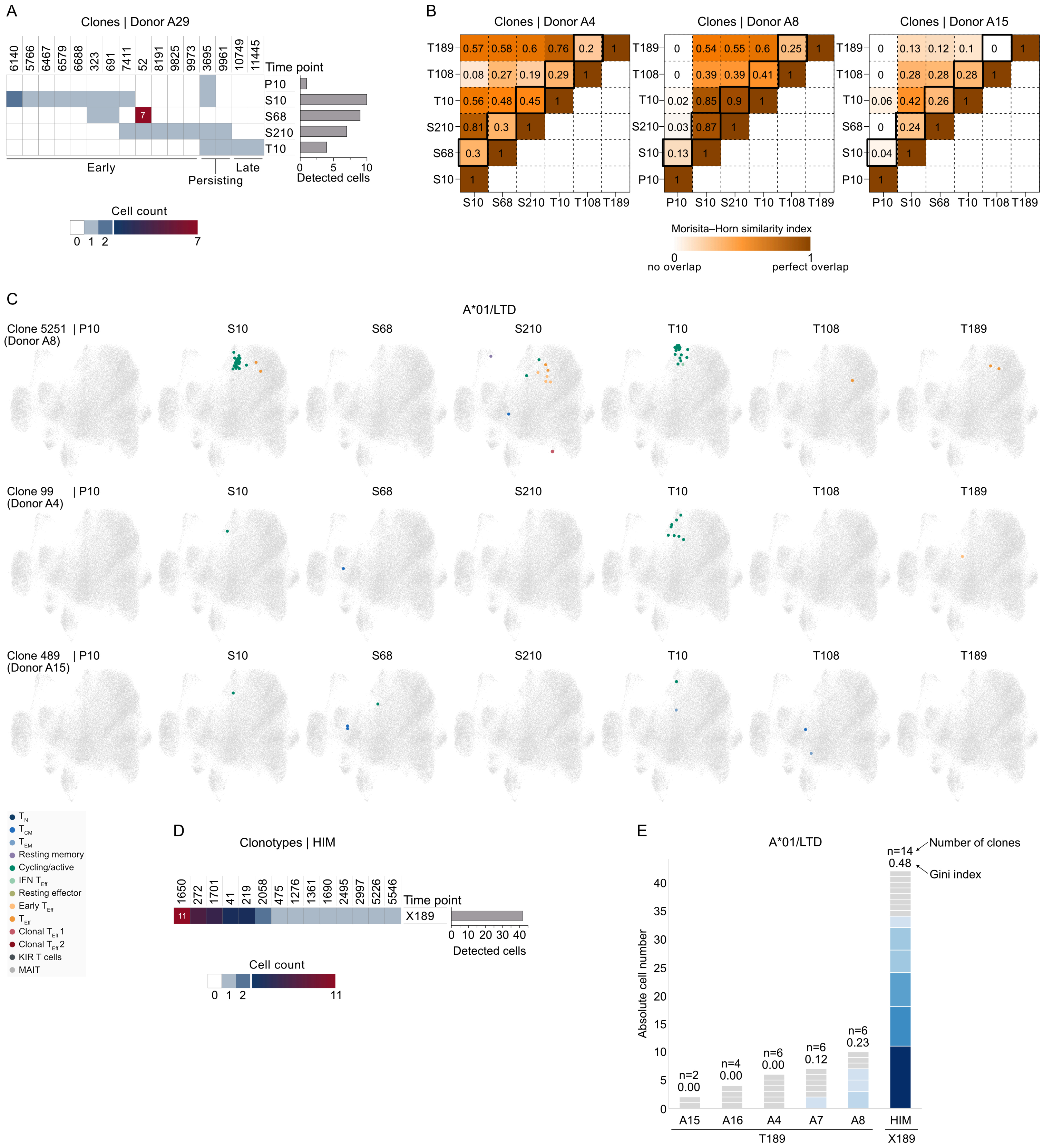


**Suppl. Fig. 7: Clonality of epitope-specific TCR repertoires. A** Detected A*02/YLQ dextramer⁺ clones across all scRNAseq experiments are shown for HLA-matched donor A29. Color gradient indicates cell counts at individual time points in days after primary (P), secondary (S) and tertiary (T) vaccination. The maximum detected cell number is indicated in the heatmap. Clones are ordered by detection pattern indicated at the bottom and by cell count at T10. Bar graphs (on the right of the heatmap) show total number of detected A*02/YLQ dextramer⁺ cells per time point. **B** Abundance-weighted similarity between A*01/LTD-specific TCR repertoires (across all scRNAseq experiments) at specific time points after vaccination for donor A4 (left), A8 (middle), and A15 (right). Color gradient indicates Morisita–Horn similarity index with exact values depicted in the heatmap. A higher index indicates higher overlap between TCR repertoires. The sampling time course is highlighted in each heatmap. **C** UMAP visualization of individual A*01/LTD dextramer⁺ clones detected at individual time points after vaccination in donor A8 (top), A4 (middle), or A15 (bottom). Colors represent cluster location of A*01/LTD dextramer⁺ cells at respective time points. Cells without the indicated epitope-specificity are shown in grey. **D** Detected A*01/LTD dextramer⁺ clones are shown for HLA-matched donor HIM 189 days after 215 reported vaccinations (X189). Color gradient indicates cell count and the maximum detected cell number is indicated in the heatmap. Clones are ordered by cell count. Bar graph (on the right of the heatmap) shows total number of detected A*01/LTD dextramer⁺ cells. **E** T cell clonality of A*01/LTD dextramer⁺ T cells of CoVa-Adapt donors 189 days after 3rd vaccination (T189) and donor HIM 189 days after 215 reported vaccinations (X189). Segments of bars indicate individual clones (grey segments = 1 cell; blue segments > 1 cell). Numbers on top of the bars represent total number of clones per donor and Gini index (high values indicate high clonality, low values indicate high evenness).

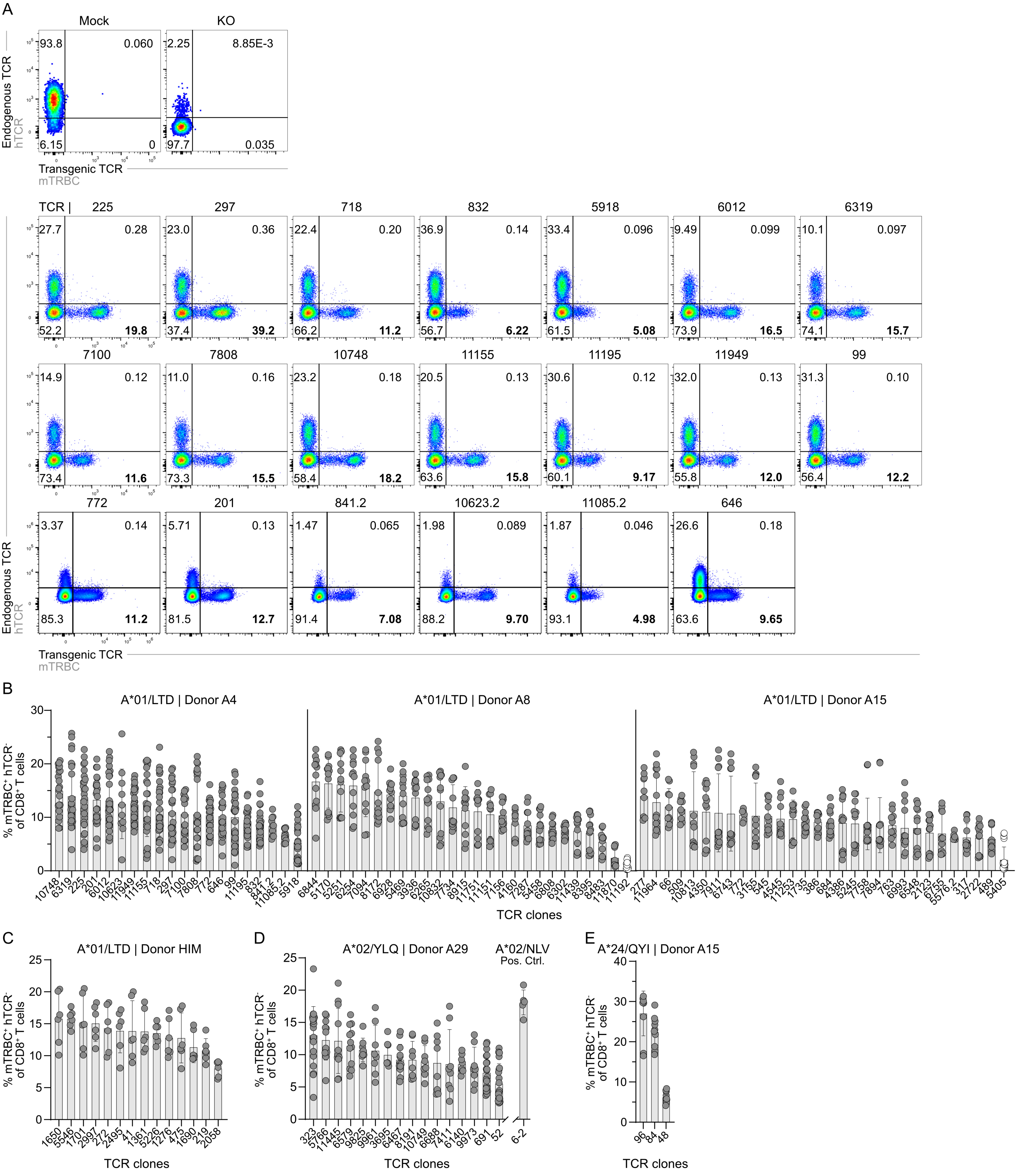


**Suppl. Fig. 8: Transgenic re-expression of epitope-specific TCRs identified by scRNAseq.** 109 SARS-CoV-2 spike epitope (A*01/LTD, A*02/YLQ, A*24/QYI) specific TCRs and one CMV (pp65, A*02/NLV) specific TCR were re-expressed in primary human T cells via CRISPR/Cas9-mediated orthotopic TCR replacement (OTR) with a murine constant region (mTRBC) to be distinguishable from the endogenous TCR (hTCR). **A** Representative flow cytometry plots four days after electroporation of the complete A*01/LTD-specific TCR repertoire of donor A4. Mock: unedited T cells, KO: T cells electroporated with RNPs targeting the endogenous *hTRAC* and *hTRBC* but no HDR template DNA (knockout). Flow cytometry plots are pre-gated on living $CD8^+$ lymphocytes. **B-E** Quantification of knockin (KI) efficiencies (protein expression) four days after electroporation of A*01/LTD-specific TCRs from CoVa-Adapt donors A4 (left, per TCR: n=10-28 technical replicates from 3-7 experiments), A8 (middle, per TCR: n=12-20; 3-5 experiments), A15 (right, per TCR: n=6-20; 2-5 experiments) (B), and donor HIM (per TCR: n=4-6; 2-3 experiments) (C); A*02/YLQ-specific TCRs from CoVa-Adapt donor A29 (per TCR: n=8-20; 2-5 experiments) and CMV (pp65, A*02/NLV) specific TCR 6-2 (n=5; 2 experiments) as positive control (Pos. Ctrl.) (D); A*24/QYI-specific TCRs from donor A15 (per TCR: n=10; 2 experiments) (E). Numbers at the bottom indicate TCR identifiers, TCRs for which a second $\alpha$– or $\beta$-chain was identified by scRNAseq and functionally tested are identified with .2 at the end. TCRs are ordered by mean KI efficiency from left to right and represented as white dots in case a TCR was classified as not re-expressed (n.r.). Data points represent technical replicates, bars with error bars show the mean +/- s.d..

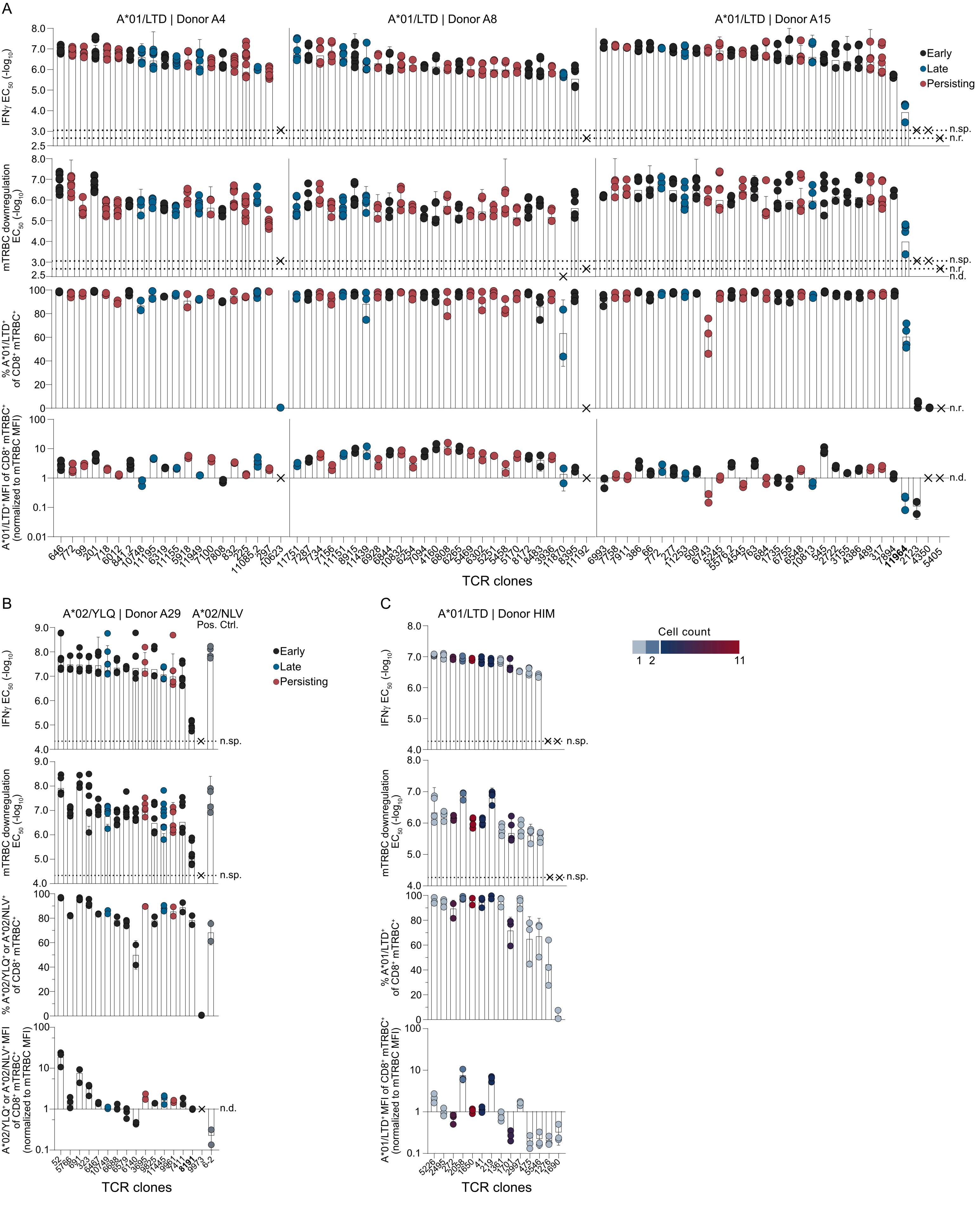

A
A*01/LTD | Donor A4
A*01/LTD | Donor A8
A*01/LTD | Donor A15
Early
Late
Persisting
IFNγ EC50 (-log10)
mTRBC downregulation EC50 (-log10)
% A*01/LTD+ of CD8+ mTRBC+
A*01/LTD+ MFI of CD8+ mTRBC+ (normalized to mTRBC MFI)
n.sp.
n.r.
n.d.
TCR clones
B
A*02/YLQ | Donor A29
A*02/NLV Pos. Ctrl.
Early
Late
Persisting
IFNγ EC50 (-log10)
mTRBC downregulation EC50 (-log10)
% A*02/YLQ+ or A*02/NLV+ of CD8+ mTRBC+
A*02/YLQ+ or A*02/NLV+ MFI of CD8+ mTRBC+ (normalized to mTRBC MFI)
n.sp.
n.d.
TCR clones
C
A*01/LTD | Donor HIM
Cell count
1
2
11
IFNγ EC50 (-log10)
mTRBC downregulation EC50 (-log10)
% A*01/LTD+ of CD8+ mTRBC+
A*01/LTD+ MFI of CD8+ mTRBC+ (normalized to mTRBC MFI)
n.sp.
TCR clones

**Suppl. Fig. 9: Functionality of epitope-specific TCR repertoires.** Quantification of peptide sensitivity ($EC_{50}$) and pHLA multimer binding of A*01/LTD-specific repertoires of donor A4, A8, and A15 (A), of a A*02/YLQ-specific TCR repertoire of donor A29 including CMV (pp65, A*02/NLV)-specific TCR (B), and A*01/LTD-specific TCR repertoire of donor HIM (C). Quantification of peptide sensitivity ($EC_{50}$) determined by dose-dependent IFNγ upregulation of living $CD8^+$ $hTCR^-$ lymphocytes (row 1), quantification of peptide sensitivity ($EC_{50}$) determined by dose-dependent transgenic TCR (mTRBC) downregulation of living $CD8^+$ $hTCR^-$ $mTRBC^+$ lymphocytes (row 2), quantification of A*01/LTD, A*02/YLQ or A*02/NLV $multimer^+$ cells of living $CD8^+$ $hTCR^-$ $mTRBC^+$ lymphocytes (row 3), quantification of A*01/LTD, A*02/YLQ or A*02/NLV multimer MFI of living $CD8^+$ $hTCR^-$ $mTRBC^+$ lymphocytes normalized to transgenic TCR expression levels (mTRBC MFI) (row 4). TCRs are ordered by increasing IFNγ $EC_{50}$ value (decreasing functionality) from left to right. Numbers at the bottom indicate TCR identifiers, TCRs for which a second α– or β-chain was identified by scRNAseq and functionally tested are identified with .2 at the end. Data points represent technical replicates, bars with error bars show the mean +/- s.d.. TCRs that were not re-expressed by OTR or did not show IFNγ upregulation above the negative control at the highest peptide concentration were labeled as not re-expressed (n.r.) or not specific (n.sp.), respectively. TCR 1276 (donor HIM) showed minimal IFNγ upregulation at the highest peptide concentration but was set to n.sp. as no $EC_{50}$ value could be calculated. mTRBC downregulation was quantified for TCRs with minimum 2% $mTRBC^+$ hTCR- cells of living $CD8^+$ lymphocytes in the unstimulated control, and labeled as not determined (n.d.) otherwise. Multimer MFI was quantified for TCRs with a minimum of five cells detected as $multimer^+$ of $CD8^+$ $hTCR^-$ $mTRBC^+$ lymphocytes, and labeled as not determined (n.d.) otherwise. **A** A*01/LTD-specific repertoires of donor A4 (left; $EC_{50}$: n=4-10 technical replicates, 2-5 experiments per TCR; % A*01/$LTD^+$: n=2-4, 2-4 experiments per TCR; A*01/LTD MFI: n=2-4, 2-4 experiments per TCR), A8 (middle; $EC_{50}$: n=4-8, 2-3 experiments per TCR; % A*01/$LTD^+$: n=2-3, 2-3 experiments per TCR; A*01/LTD MFI: n=2, 2 experiments per TCR ), and A15 (right; IFNγ $EC_{50}$: n=4-6, 2-3 experiments per TCR; mTRBC $EC_{50}$: n=2-6, 1-3 experiments per TCR; % A*01/$LTD^+$: n=2-4, 2-4 experiments per TCR; A*01/LTD MFI: n= 2-3, 2-3 experiments per TCR). TCRs are colored based on their detection pattern (see legend on top right) and TCR 11964 with ultra-low avidity ($EC_{50}$ of $-\log_{10}$ 3.94M) is highlighted in bold. **B** A*02/YLQ-specific repertoire of donor A29 (IFNγ $EC_{50}$: n=4-6, 2-3 experiments per TCR; mTRBC $EC_{50}$: n= 3-8, 2-4 experiments; % A*02/$YLQ^+$: n=2-3, 2-3 experiments per TCR; A*01/LTD MFI: n=2-3, 2-3 experiments per TCR) including CMV (pp65, A*02/NLV)-specific TCR 6-2 as positive control (Pos. Ctrl.; $EC_{50}$: n=4, 2 experiments per TCR; % A*02/$NLV^+$: n=2, 2 experiments per TCR; A*02/NLV MFI: n=2, 2 experiments per TCR). TCRs are colored based on their detection pattern (see legend on the right) and TCR 8191 with ultra-low avidity ($EC_{50}$ of $-\log_{10}$ 4.91M) is highlighted in bold. **C** A*01/LTD-specific repertoire of donor HIM ($EC_{50}$: n=4, 2 experiments per TCR; % A*01/$LTD^+$: n=3, 3 experiments per TCR; A*01/LTD MFI: n=3, 3 experiments per TCR). TCRs are colored based on degree of clonal expansion in scRNAseq dataset (see legend on the right).

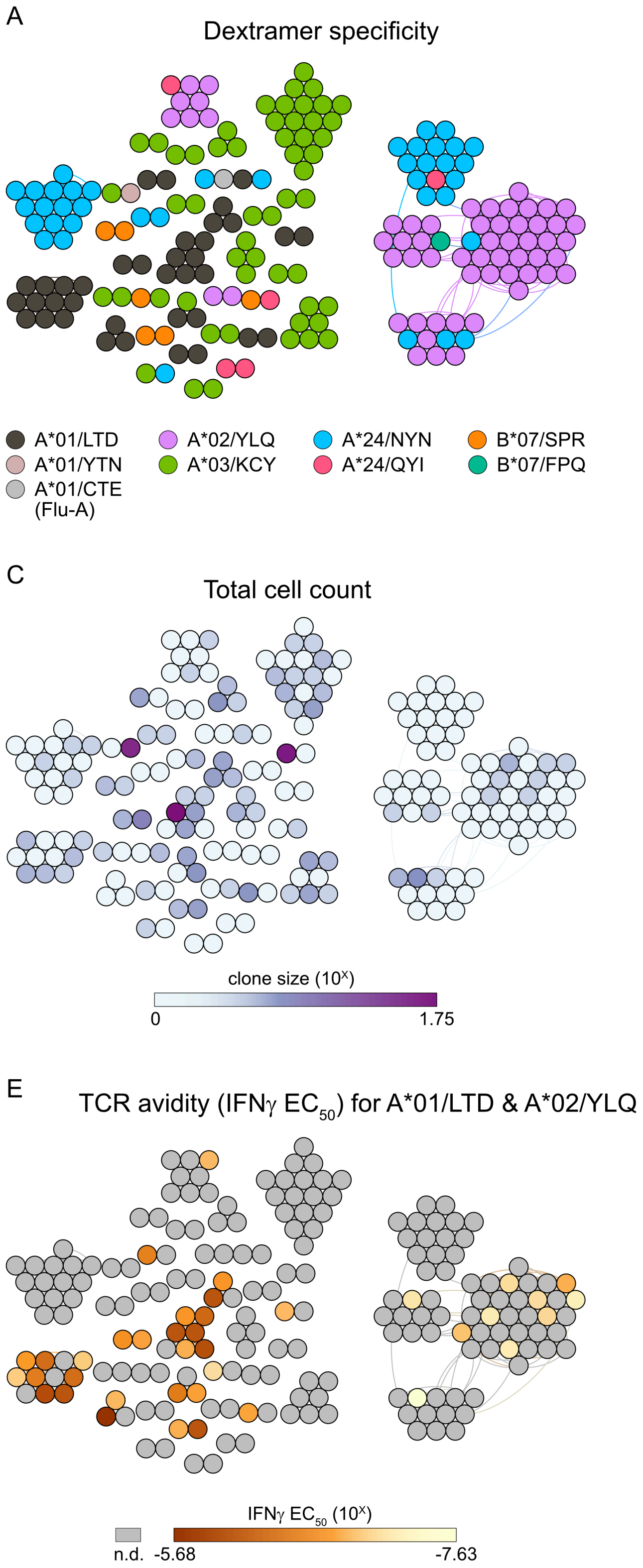


**Suppl. Fig. 10: TCR similarity clusters of epitope-specific TCRs.** Similarity network of TCRs identified as dextramer$^{+}$ in scRNAseq (all SARS-CoV-2 spike, Flu-A (A*01/CTE), and EBV (B*08/FLR, B*08/RAK) epitope specificities). Each vertex on a similarity network represents a unique paired αβTCR clonotype, and edges connect vertices with <120 TCRdist units. Only clusters with at least two clone members are shown. No TCR clone with ultra-low avidity ($EC_{50}$ values < $-\log_{10}$ 5M) or tested as not specific became part of one of the clusters based on these criteria. Colors indicate epitope specificity (**A**), detection pattern (**B**), total cell count over all acquired time points pooled (**C**), cell count at T10 (**D**), TCR avidity determined by dose-dependent IFNγ upregulation of living $CD8^{+}$ $hTCR^{-}$ lymphocytes, clones without determined TCR avidity are labeled in grey for not determined (n.d.) (**E**), or differentiation degree indicated by diffusion pseudotime (high pseudotime, high differentiation degree) (**F**).

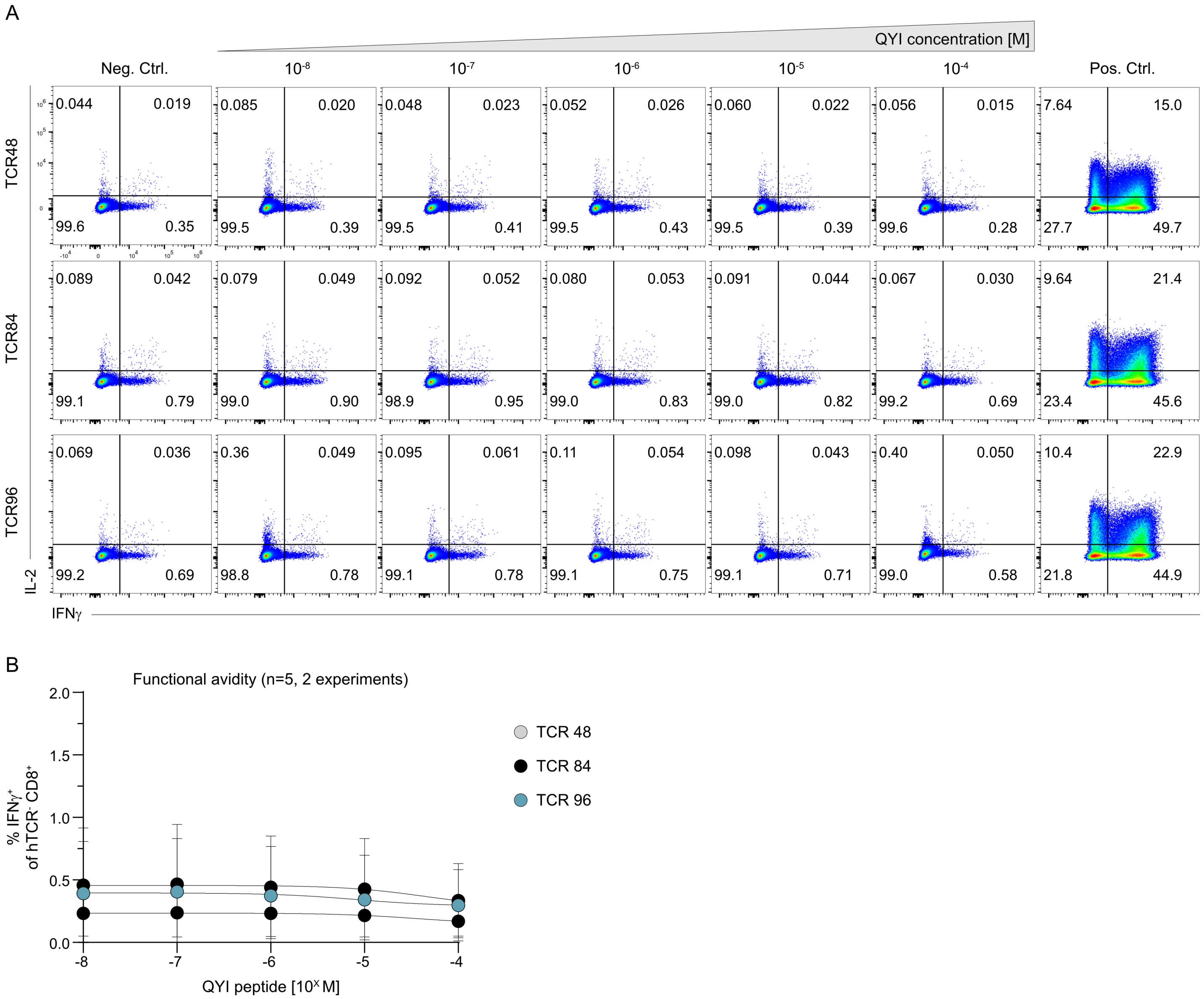


**Suppl. Fig. 11: Functionality of A*24/QYI dextramer$^+$ TCRs.** A*24/QYI dextramer$^+$ TCRs were transgenically re-expressed in primary human T cells by CRISPR/Cas9-mediated orthotopic TCR replacement (OTR) and tested for peptide sensitivity ($EC_{50}$) twelve days after OTR. TCR-engineered T cells were co-cultured with APCs loaded with QYI peptide ranging from $10^{-8}$ to $10^{-4}$ M in a ratio of 1:1 for 4h. **A** Primary flow cytometry data are pre-gated on living CD8$^+$ hTCR$^-$ lymphocytes. Negative control (Neg. Ctrl) = solvent, positive control (Pos. Ctrl) = PMA/ionomycin. **B** Quantification of dose-dependent IFNγ upregulation of living CD8$^+$ hTCR$^-$ lymphocytes (n=5 technical replicates from 2 experiments). Dots with error bars show the mean +/- s.d. and are colored according to TCR clonotype.

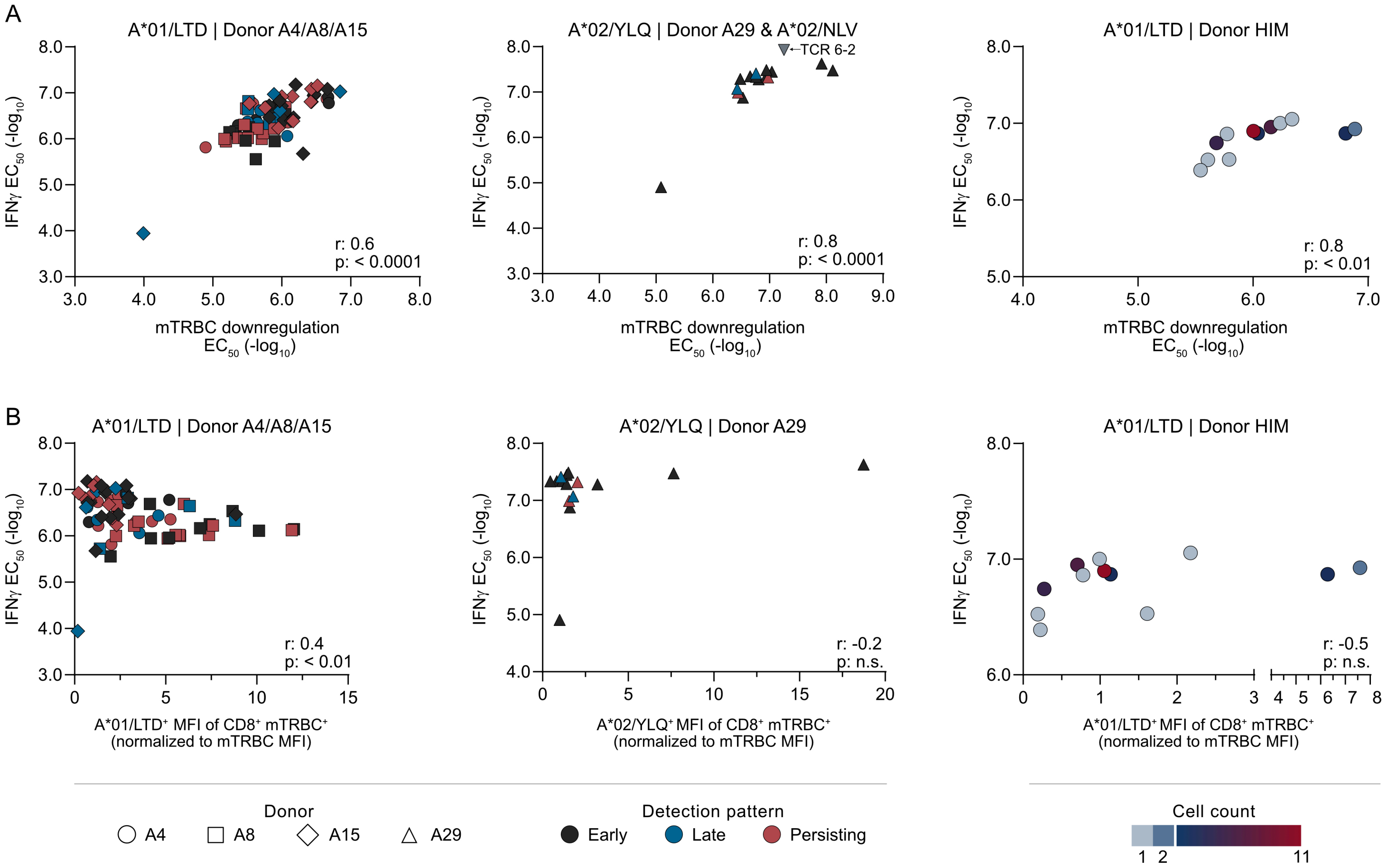


**Suppl. Fig. 12: Correlation of TCR avidity parameters. A** Correlation of peptide sensitivity ($EC_{50}$) determined by dose-dependent IFNγ upregulation of living CD8⁺ hTCR⁻ lymphocytes (IFNγ $EC_{50}$) and determined by dose-dependent transgenic TCR (mTRBC) downregulation of living CD8⁺ hTCR⁻ mTRBC⁺ lymphocytes (mTRBC downregulation $EC_{50}$) for A*01/LTD-specific TCRs of donor A4, A8, and A15 (left, n=70 TCR clones), A*02/YLQ-specific TCRs from donor A29 (n=15 TCR clones) including CMV (pp65, A*02/NLV)-specific TCR 6-2 as positive control (middle), and A*01/LTD-specific TCRs of donor HIM (n=12 TCR clones) (right). **B** Correlation of peptide sensitivity (IFNγ $EC_{50}$) as in (A) and A*01/LTD or A*02/YLQ multimer MFI of living CD8⁺ hTCR⁻ mTRBC⁺ lymphocytes normalized to transgenic TCR expression levels (mTRBC MFI) for A*01/LTD-specific TCRs of donor A4, A8, and A15 (left, n=71 TCR clones), A*02/YLQ-specific TCRs from donor A29 (middle, n=15 TCR clones), and A*01/LTD-specific TCRs of donor HIM (right, n=12 TCR clones). Data points represent individual clones, symbols represent individual donors, and color represents detection pattern (donors A4, A8, A15, A29) or cell count (HIM) of each clone (see legend on bottom left and right). Correlation was analyzed by non-parametric spearman correlation and indicated in the figure.

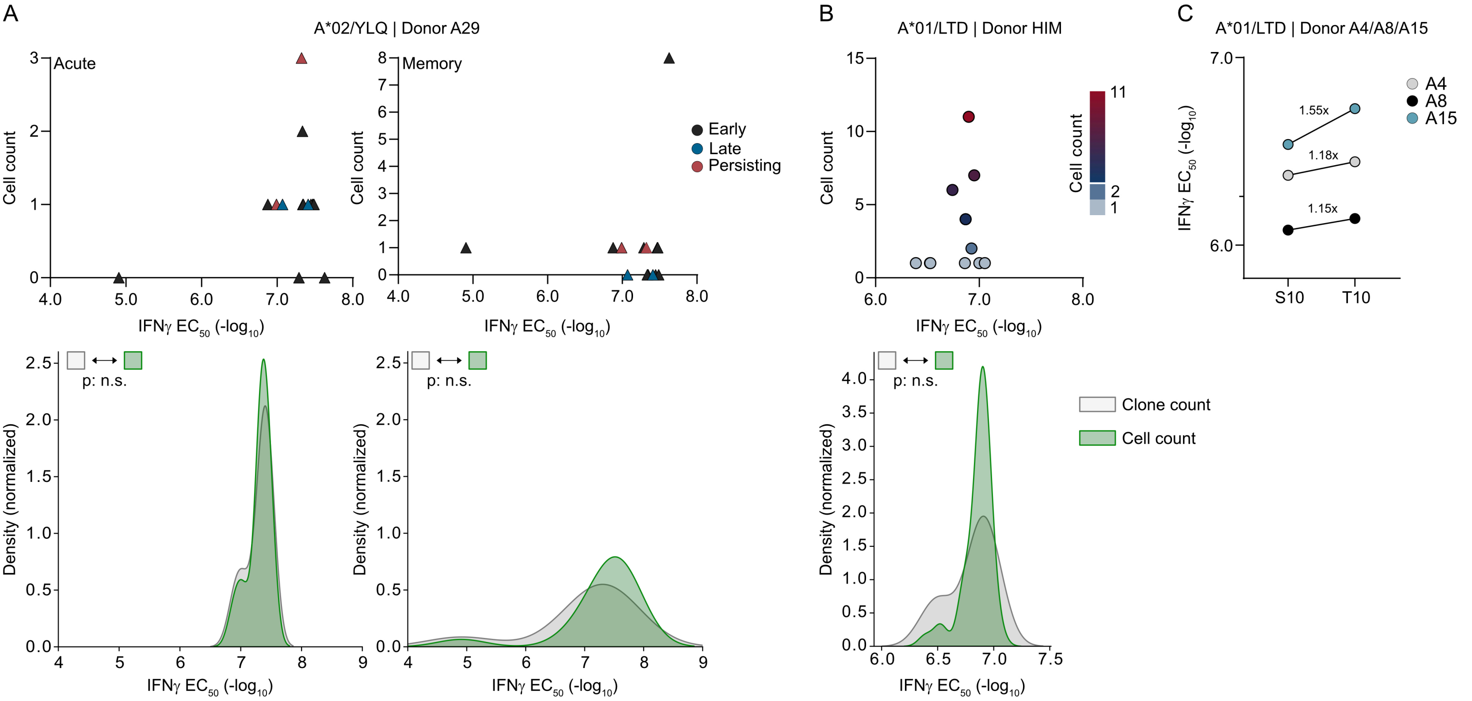


**Suppl. Fig. 13: Correlation of TCR functionality and clonal expansion. A** Correlation of $EC_{50}$ values (determined by dose-dependent IFNγ upregulation) with cell counts of A*02/YLQ-specific TCRs (donor A29) in acute (P10/S10/T10) and memory (S68/S210) phase after vaccination (top). Data points represent individual TCR clones and color represents detection pattern of each clone (see legend on the right). Clone count (grey) and cell count (green) distribution for A*02/YLQ-specific clones from repertoire of donor A29 (bottom). Clone and cell count is depicted for acute (P10/S10/T10) and memory (S68/S210) phase after vaccination. Statistical comparison of clone and cell count distribution was performed by Kolmogorow-Smirnoff test, n.s., not significant. **B** Correlation of $EC_{50}$ values (determined by dose-dependent IFNγ upregulation) with cell counts of A*01/LTD-specific TCRs (donor HIM) 189 days after 215 reported vaccinations (X189) (top). Data points represent individual TCR clones and color represents cell count (see legend on the right). Clone count (grey) and cell count (green) distribution for A*01/LTD-specific clones from repertoire of donor HIM (bottom). Statistical comparison of clone and cell count distribution was performed by Kolmogorow-Smirnoff test, n.s., not significant. **C** TCR avidity (determined by dose-dependent IFNγ upregulation) was determined for the complete A*01/LTD-specific population detected at S10 or T10 for donors A4, A8, and A15. Population avidity was calculated weighing in clonal expansion (see Methods). Colored dots represent donor-specific population avidity. Fold increase in population avidity is indicated for each individual donor.

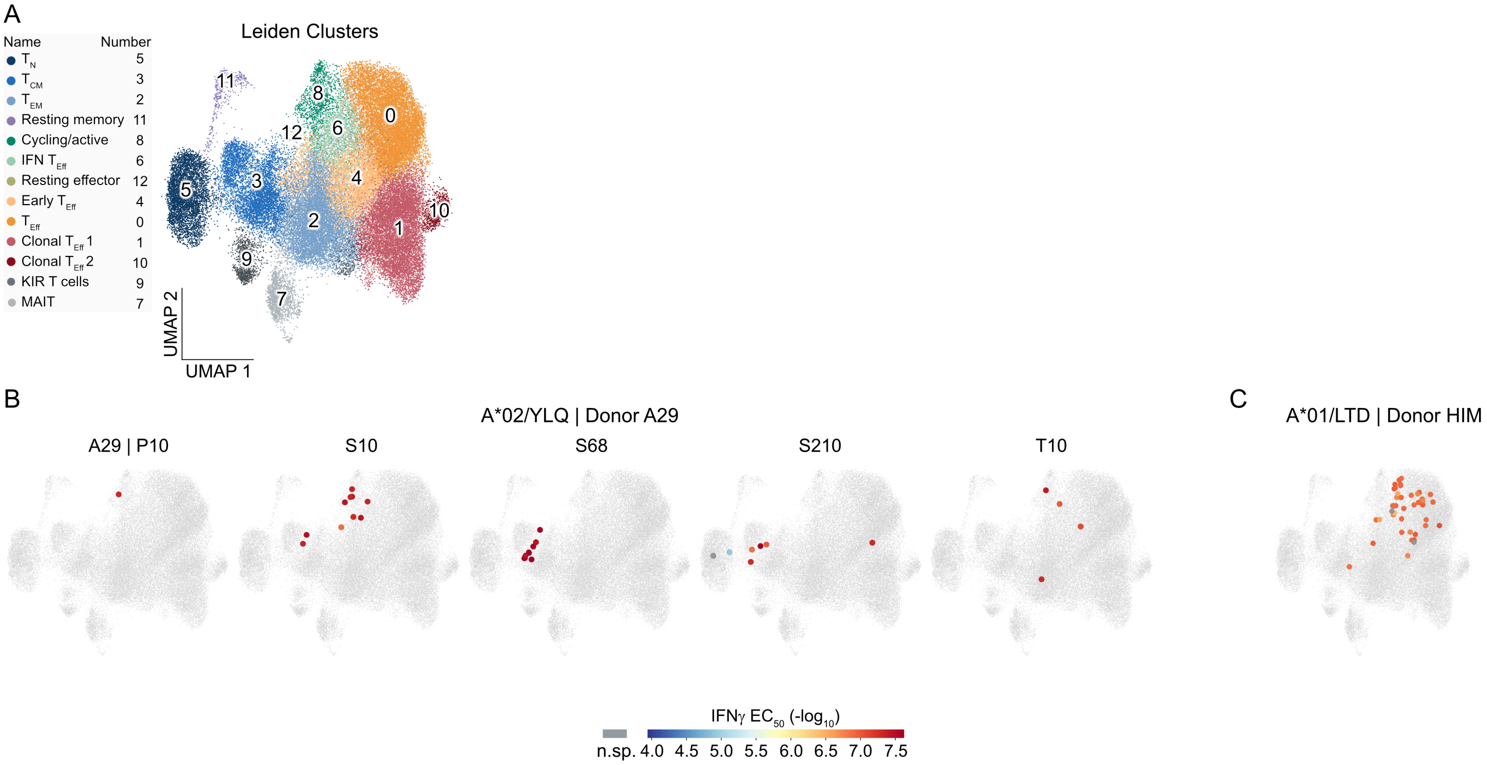


**Suppl. Fig. 14: Longitudinal phenotypes of functionally characterized epitope-specific TCRs. A** UMAP with Leiden clusters (cluster numbers in UMAP, names and numbers on the left) of CD8$^{+}$ T cells enriched for dextramer-binding (Suppl. Fig. 2A) from three independent scRNAseq experiments (n = 53,907 cells). **B** Visualization of cells belonging to all functionally characterized A*02/YLQ dextramer$^{+}$ clones from donor A29 on UMAP at individual time points in days after primary (P), secondary (S) and tertiary (T) vaccination (colored circled large dots; color gradient indicates IFNγ $EC_{50}$ values). Cells of clones that did not show IFNγ upregulation above the negative control at the highest peptide concentration were labeled as not specific (n.sp.) and are shown in dark grey and circled large dots. TCR 1276 (donor HIM) showed minimal IFNγ upregulation at the highest peptide concentration but was set to n.sp. as no $EC_{50}$ value could be calculated. Not functionally characterized cells are shown in light grey and uncircled small dots. **C** Visualization of functionally characterized A*01/LTD dextramer$^{+}$ clones from donor HIM 189 days after 215 reported vaccinations (X189).

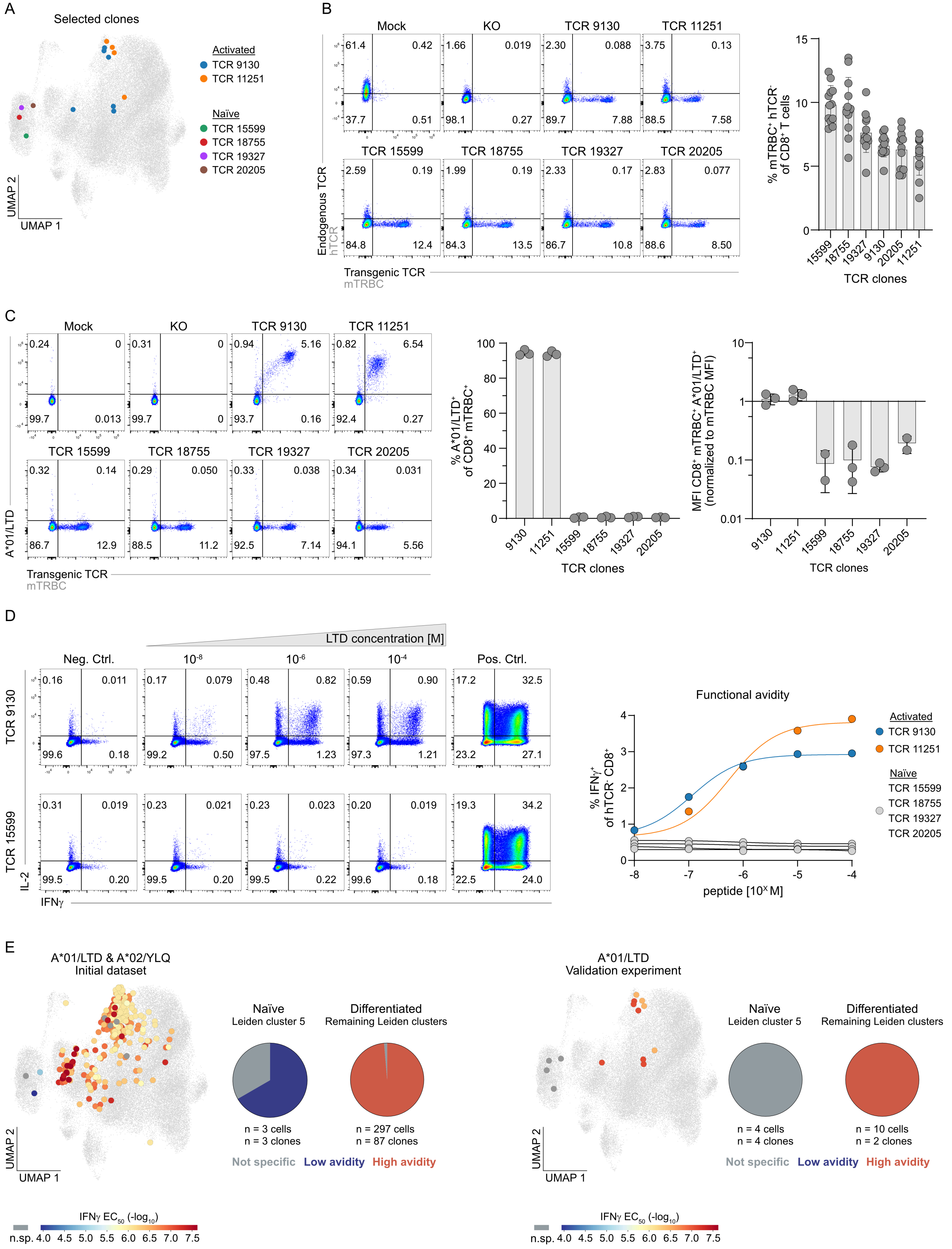

A
Selected clones
Activated
TCR 9130
TCR 11251
Naïve
TCR 15599
TCR 18755
TCR 19327
TCR 20205
UMAP 2
UMAP 1
B
Mock
KO
TCR 9130
TCR 11251
TCR 15599
TCR 18755
TCR 19327
TCR 20205
Endogenous TCR
hTCR
Transgenic TCR
mTRBC
% mTRBC⁺ hTCR⁻ of CD8⁺ T cells
TCR clones
C
A*01/LTD
% A*01/LTD⁺ of CD8⁺ mTRBC⁺
MFI CD8⁺ mTRBC⁺ A*01/LTD⁺ (normalized to mTRBC MFI)
D
LTD concentration [M]
Neg. Ctrl.
Pos. Ctrl.
IL-2
IFNγ
Functional avidity
% IFNγ⁺ of hTCR⁻ CD8⁺
peptide [10^X M]
E
A*01/LTD & A*02/YLQ
Initial dataset
Naïve
Leiden cluster 5
Differentiated
Remaining Leiden clusters
n = 3 cells
n = 3 clones
n = 297 cells
n = 87 clones
Not specific
Low avidity
High avidity
IFNγ EC50 (-log10)
n.sp.
A*01/LTD
Validation experiment
n = 4 cells
n = 4 clones
n = 10 cells
n = 2 clones

**Suppl. Fig. 15: Validation experiment for A*01/LTD TCR functionality. A** UMAP visualization of newly ordered TCRs belonging to A*01/LTD dextramer$^+$ clones with recruitment into cycling/active cluster 8 (activated) or with solely a naïve (cluster 5) phenotype. **B** Six A*01/LTD dextramer$^+$ TCRs were re-expressed in primary human T cells via by CRISPR/Cas9-mediated orthotopic TCR replacement (OTR) with a murine constant region (mTRBC) to be distinguishable from the endogenous TCR (hTCR). Representative flow cytometry plots (left) four days after electroporation; Mock: unedited T cells, KO: T cells electroporated with RNPs targeting the endogenous *hTRAC* and *hTRBC* but no HDR template DNA (knockout). Flow cytometry plots are pre-gated on living CD8$^+$ lymphocytes. Quantification (right) of knockin (KI) efficiencies (protein expression) four days after electroporation (n=12 technical replicates from 3 experiments per TCR). TCRs are ordered by KI efficiency from left to right. **C** On day 11 after OTR, TCR-engineered T cells were stained with A*01/LTD multimer. Primary data are pre-gated on living CD8$^+$ lymphocytes (left). Quantification of A*01/LTD multimer$^+$ cells of living CD8$^+$ hTCR$^-$ mTRBC$^+$ lymphocytes (middle) and quantification of A*01/LTD multimer MFI of living CD8$^+$ hTCR$^-$ mTRBC$^+$ lymphocytes normalized to transgenic TCR expression levels (mTRBC MFI) (right). **C** A*01/LTD dextramer$^+$ TCRs were tested for peptide sensitivity ($EC_{50}$) at day 12 after OTR. TCR-engineered T cells were co-cultured with APCs loaded with LTD peptide ranging from $10^{-8}$ to $10^{-4}$ M in a ratio of 1:1 for 4h. Primary data (left) are pre-gated on living CD8$^+$ hTCR$^-$ lymphocytes. Negative control (Neg. Ctrl) = solvent, positive control (Pos. Ctrl) = PMA/ionomycin. Quantification (right) of dose-dependent IFNγ upregulation of living CD8$^+$ hTCR$^-$ lymphocytes for one representative experiment. Dots are colored according to TCR for activated clones and grey for not specific (n.sp.) clones from naïve cluster 5. **E** UMAP visualization of all initially re-expressed and functionally characterized A*01/LTD (donor A4, A8, A15) and A*02/YLQ dextramer$^+$ (A29) TCRs for all time points pooled (left) and newly ordered A*01/LTD dextramer$^+$ TCRs (right) (colored circled large dots; color gradient indicates IFNγ $EC_{50}$ values). Cells of clones that did not show IFNγ upregulation above the negative control at the highest peptide concentration were labeled as not specific (n.sp.) and are shown in dark grey and circled large dots. Not functionally characterized cells are shown in light grey and uncircled small dots. TCR clones were separated into not specific, low avidity (IFNγ $EC_{50}$ values < $-\log_{10}$ 5M), and high avidity (IFNγ $EC_{50}$ values > $-\log_{10}$ 5M) clones. Pie charts represent fractional distribution of cells from these three clone groups within the naïve cluster 5 and all other Leiden clusters. Color represents clone group, total cell and clone number is indicated under individual pie charts.

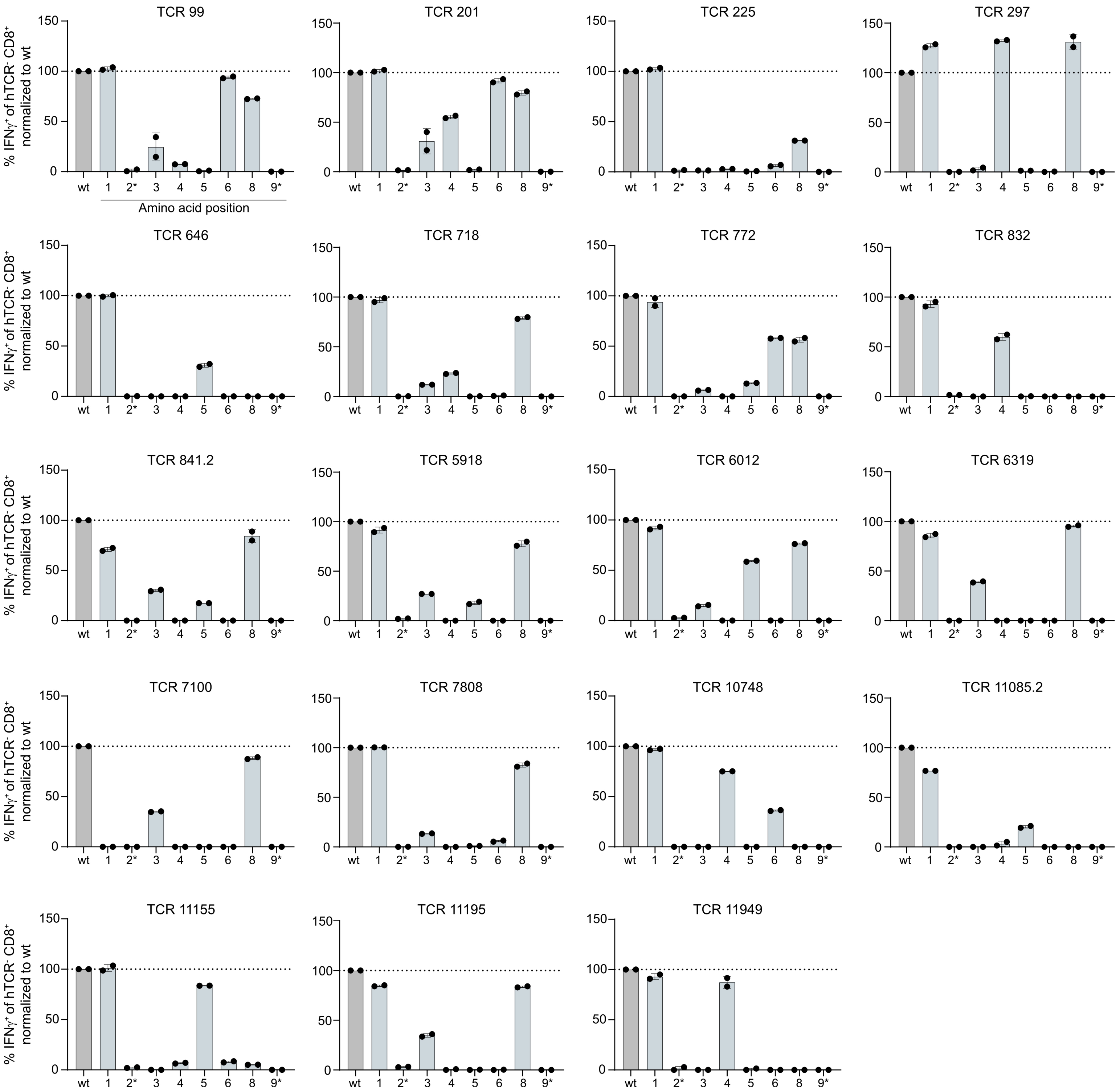


**Suppl. Fig. 16: Reactivity landscape of A*01/LTD specific TCRs against altered peptide ligands.** Engineered T cells expressing A*01/LTD-specific TCRs from donor A4 were co-cultured with APCs loaded with $10^{-5}$ M of wild type (wt) LTD or altered peptide ligands (APLs) in a ratio of 1:1 for 4h. Quantification of IFNγ upregulation normalized to stimulation with wt LTD (100%, dotted line) for each individual TCR and each APL with one position at a time being mutated to alanine. The mutated amino acid position is indicated with numbers at the bottom of each graph with anchor positions highlighted with asterisks.
